# Large-scale single-cell analysis reveals critical immune characteristics of COVID-19 patients

**DOI:** 10.1101/2020.10.29.360479

**Authors:** Xianwen Ren, Wen Wen, Xiaoying Fan, Wenhong Hou, Bin Su, Pengfei Cai, Jiesheng Li, Yang Liu, Fei Tang, Fan Zhang, Yu Yang, Jiangping He, Wenji Ma, Jingjing He, Pingping Wang, Qiqi Cao, Fangjin Chen, Yuqing Chen, Xuelian Cheng, Guohong Deng, Xilong Deng, Wenyu Ding, Yingmei Feng, Rui Gan, Chuang Guo, Weiqiang Guo, Shuai He, Chen Jiang, Juanran Liang, Yi-min Li, Jun Lin, Yun Ling, Haofei Liu, Jianwei Liu, Nianping Liu, Yang Liu, Meng Luo, Qiang Ma, Qibing Song, Wujianan Sun, GaoXiang Wang, Feng Wang, Ying Wang, Xiaofeng Wen, Qian Wu, Gang Xu, Xiaowei Xie, Xinxin Xiong, Xudong Xing, Hao Xu, Chonghai Yin, Dongdong Yu, Kezhuo Yu, Jin Yuan, Biao Zhang, Tong Zhang, Jincun Zhao, Peidong Zhao, Jianfeng Zhou, Wei Zhou, Sujuan Zhong, Xiaosong Zhong, Shuye Zhang, Lin Zhu, Ping Zhu, Bin Zou, Jiahua Zou, Zengtao Zuo, Fan Bai, Xi Huang, Xiuwu Bian, Penghui Zhou, Qinghua Jiang, Zhiwei Huang, Jin-Xin Bei, Lai Wei, Xindong Liu, Tao Cheng, Xiangpan Li, Pingsen Zhao, Fu-Sheng Wang, Hongyang Wang, Bing Su, Zheng Zhang, Kun Qu, Xiaoqun Wang, Jiekai Chen, Ronghua Jin, Zemin Zhang

## Abstract

Dysfunctional immune response in the COVID-19 patients is a recurrent theme impacting symptoms and mortality, yet the detailed understanding of pertinent immune cells is not complete. We applied single-cell RNA sequencing to 284 samples from 205 COVID-19 patients and controls to create a comprehensive immune landscape. Lymphopenia and active T and B cell responses were found to coexist and associated with age, sex and their interactions with COVID-19. Diverse epithelial and immune cell types were observed to be virus-positive and showed dramatic transcriptomic changes. Elevation of ANXA1 and S100A9 in virus-positive squamous epithelial cells may enable the initiation of neutrophil and macrophage responses via the ANXA1-FPR1 and S100A8/9-TLR4 axes. Systemic upregulation of S100A8/A9, mainly by megakaryocytes and monocytes in the peripheral blood, may contribute to the cytokine storms frequently observed in severe patients. Our data provide a rich resource for understanding the pathogenesis and designing effective therapeutic strategies for COVID-19.

**HIGHLIGHTS:**

- Large-scale scRNA-seq analysis depicts the immune landscape of COVID-19
- Lymphopenia and active T and B cell responses coexist and are shaped by age and sex
- SARS-CoV-2 infects diverse epithelial and immune cells, inducing distinct responses
- Cytokine storms with systemic S100A8/A9 are associated with COVID-19 severity

## INTRODUCTION

The coronavirus disease 2019 (COVID-19) is an ongoing pandemic infectious disease, caused with the severe acute respiratory syndrome coronavirus 2 (SARS-CoV-2). Currently, it has caused with around 29 million infections and close to 1 million deaths according to the statistics of World Health Organization until September 15, 2020, with the fatality rate as high as ~10% in specific regions. Although many COVID-19 patients experience asymptomatic, mild or moderate symptoms, some patients progress to severe conditions and even death. It is thus of paramount importance to understand the disease mechanisms and the underlying factors associated with vulnerabilities, which are critical for controlling the pandemic and alleviating the global crisis. It is also critical to systematically investigate differences between clinical presentations (mild/moderate and severe), or between treatment outcomes (disease progression and convalescence) of patients, as they can provide important guidance to the development of effective therapeutics and vaccines.

Multiple studies have suggested the alterations of immune responses as one of the key mechanisms for severe symptoms (Guo et al., 2020a; Schulte-Schrepping et al., 2020; Silvin et al., 2020a; Wen et al., 2020; Zhang et al., 2020a; Zhang et al., 2020b). Patients with severe COVID-19 might have a cytokine storm syndrome accompanying the hyper-inflammatory response, which is a major cause of disease severity and death (Cao, 2020; Del Valle et al., 2020; Huang et al., 2020; Liao et al., 2020; Mehta et al., 2020; Zhou et al., 2020a). During the inflammatory responding process, pathogenic T cells and inflammatory monocytes produced inflammatory cytokines (Zhou et al., 2020b) such as G-CSF (Costela-Ruiz et al., 2020; Du et al., 2020), TNF-α (Jamilloux et al., 2020; Vabret et al., 2020), IL-6 and IL-1 (Abbasifard and Khorramdelazad, 2020; Costela-Ruiz et al., 2020; Del Valle et al., 2020; Liao et al., 2020; Mehta et al., 2020; Yang et al., 2020), and drove downstream hyper-inflammation. In contrast, some studies have argued against the presence of cytokine storms (Kox et al., 2020; Wilk et al., 2020). Thus, the detailed immune responses in COVID-19 patients with SARS-CoV-2 infection need to be more thoroughly investigated.

Single-cell RNA sequencing (scRNA-seq) is powerful at dissecting the immune responses under various conditions at the finest resolution, and has been applied to COVID-19 studies on limited scales (Chua et al., 2020; Guo et al., 2020a; He et al., 2020; Liao et al., 2020; Wilk et al., 2020; Xie et al., 2020). While the current single cell studies of COVID-19 have provided certain details of the cellular and molecular changes of patients after SARS-CoV-2 infection and even during convalescence (Mathew et al., 2020a), the small sample sizes of such studies have raised concerns over the robustness and the generalization of such findings. Here we applied scRNA-seq to a large cohort with 205 individuals, including hospitalized COVID-19 patients with moderate or severe disease, and patients in the convalescent stage, as well as healthy controls. With high-quality transcriptomics data of ~1.5 million single cells, we reveal that SARS-CoV-2 could infect a wider range of cell types than previous understanding, and induce distinct phenotypic changes in those infected cells. Such heterogeneity of SARS-CoV-2 infection has important immunological implications as such cells exhibit distinct interaction potentials with innate and adaptive immune cells. We also observed critical changes in the peripheral blood discriminating mild/moderate from severe COVID-19 patients in the disease progression or convalescence stages, and found their association with patient sex and age. Further, our large cohort analysis provides a unique opportunity to reveal the characteristics of cytokine storms in patients, and to further illustrate the cell subpopulations that might contribute to the inflammatory responses and the hyper-inflammatory genomic signatures under SARS-CoV-2 infection. Our findings may have important implications to the research, treatment, control and prevention of COVID-19.

## RESULTS

### Integrated analysis of the COVID-19 scRNA-seq data

To systematically characterize the immune properties at single-cell resolution in the COVID-19 patients, we formed a Single Cell Consortium for COVID-19 in China (SC4), which consisted of researchers from 36 research institutes or hospitals from different regions of China. Members of SC4 contributed COVID-19 related scRNA-seq data, mostly still unpublished, for a total of 205 individuals, including 25 patients with mild/moderate symptoms, 63 hospitalized patients with severe symptoms, and 92 recovered convalescent persons, as well as 25 healthy controls (Figure 1A and Table S1). While most previous studies did not discriminate whether convalescent individuals recovered from mild/moderate or severe symptoms, we divided the convalescent group into two subgroups, 54 recovered from mild/moderate symptoms and 38 recovered from severe symptoms, to investigate the effects of disease severity on the immune status of recovered individuals. This cohort covered a wide age range (from 6 to 92 years old), with the mild/moderate and severe groups having significant age differences (Figure S1A), consistent with the epidemiological observations that aged patients are prone to severe symptoms (Guo et al., 2020a; Hadjadj et al., 2020; Silvin et al., 2020b; Wilk et al., 2020; Yu et al., 2020). Additionally, no significant difference was noted in the sex composition between the mild/moderate and severe groups (Figure S1B).

**Figure 1.**
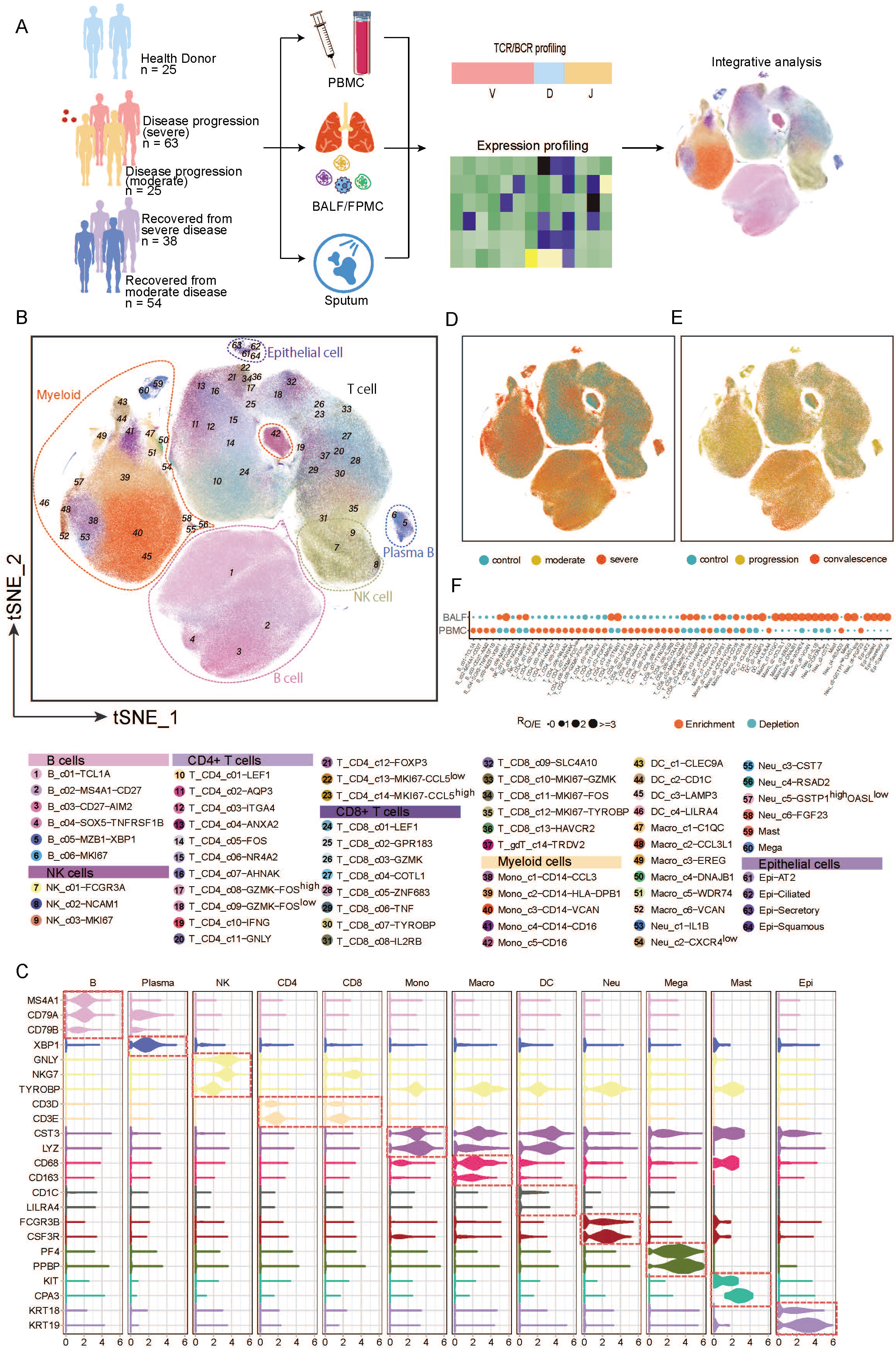
Single-cell atlas of multiple tissue types from healthy individuals and COVID-19 patients. (A) A flowchart depicting the overall design of the study. (B) t-Stochastic Neighborhood Embedding (t-SNE) representations of integrated single-cell transcriptomes of 1,462,702 cells derived from our healthy controls and COVID-19 patients. Cells are colour-coded by 64 cell subsets from 6 major cell types. (C) Violin plots of selected marker genes (rows) for major cell subpopulations (columns) ordered by cell lineage relationships. NK, natural killer cells; Mono, monocyte; Macro, macrophage; DC, dendritic cells; Neu, neutrophil; Mega, megakaryocyte; Epi, epithelial cells. (D-E) t-SNE representations of integrated single-cell transcriptomes of 1,462,702 cells coloured by disease symptoms (D) and disease progression stages (E). (F) Tissue preference of each cluster estimated by Ro/e. Ro/e denotes the ratio of observed to expected cell number. See also Figure S1.

A total of 284 samples were collected for scRNA-seq, of which 249 were from peripheral blood mononuclear cells (PBMCs) and 35 from the respiratory system, which was further composed of 12 bronchoalveolar lavage fluid (BALF) samples, 22 sputum samples, and 1 sample for pleural fluid mononuclear cells. Some patients had multiple samples collected, including seven patients with matching BALF and PBMC. Most samples were subjected to scRNA-seq based on the 10X Genomics 5’ sequencing platform to generate both the gene expression and T cell receptor (TCR) or B cell receptor (BCR) data. The scRNA-seq raw data were analyzed by a unified analysis pipeline, including the kallisto and bustools programs (Bray et al., 2016; Melsted et al., 2019), to obtain the gene expression data of individual cells and by the CellRanger program to obtain TCR and BCR sequences.

We applied a common set of stringent quality control criteria to ensure that the selected data were from single and live cells and that their transcriptomic phenotypes were comprehensively characterized. A total of 1,462,702 high-quality single cells were ultimately obtained, with an average of 4,835 unique molecular identifiers (UMIs), representing 1,587 genes (Figures S1D and S1E). With the large-scale of data, we obtained 64 cell clusters, covering diverse epithelial cells in the respiratory system, megakaryocytes, mast cells, myeloid cells, and NK/T/B cells (Figure 1B). Such an information-rich resource (available at http://covid19.cancer-pku.cn/ for quick browsing) enabled accurate annotation and analysis of these cell clusters at different resolutions (Figure 1C, Figure S1F-J and Table S2), which allow the elucidation of potential molecular and cellular mechanisms underlying the pathogenesis of SARS-CoV-2 infection and differences of human immune responses for patients with distinct symptoms.

Notable differences could be observed in the immune compositions of healthy controls and COVID-19 patients with mild/moderate or severe symptoms (Figure 1D) or between the disease progression stages and convalescence (Figure 1E) based on the t-distributed stochastic neighbor embedding (t-SNE) projection. The tissue preference of each cluster was illustrated based on the ratio of observed to randomly expected cell numbers (Ro/e, Figure 1F), partially reflecting the validity of cell clustering. Notably, various clusters of proliferating CD8+ and CD4+ T, and plasma B cells were more enriched in BALF than PBMCs, indicating activated adaptive immune responses in the lung (Figure 1F).

We first analyzed the compositional changes of the broad categories of immune cells for PBMCs in different COVID-19 patient groups. Notably, the percentages of megakaryocytes and monocytes in PBMCs were elevated, particularly in severe COVID-19 patients during the disease progression stage (Figure 2A) (Guo et al., 2020a; Zhang et al., 2020b). While NK cells did not show significant changes among the different patient groups (Figure 2A), B cells were significantly increased in severe COVID-19 patients (Figure 2A)(Guo et al., 2020a; Mathew et al., 2020b; Zhang et al., 2020b). By contrast, T cells and DCs were decreased in severe COVID-19 patients (Figure 2A). These findings are consistent with previous reports that lymphopenia is frequently observed in COVID-19 patients and that impaired adaptive immunity may occur (Gao et al., 2020; Giamarellos-Bourboulis et al., 2020; Kuri-Cervantes et al., 2020; Ni et al., 2020a; Yu et al., 2020).

**Figure 2.**
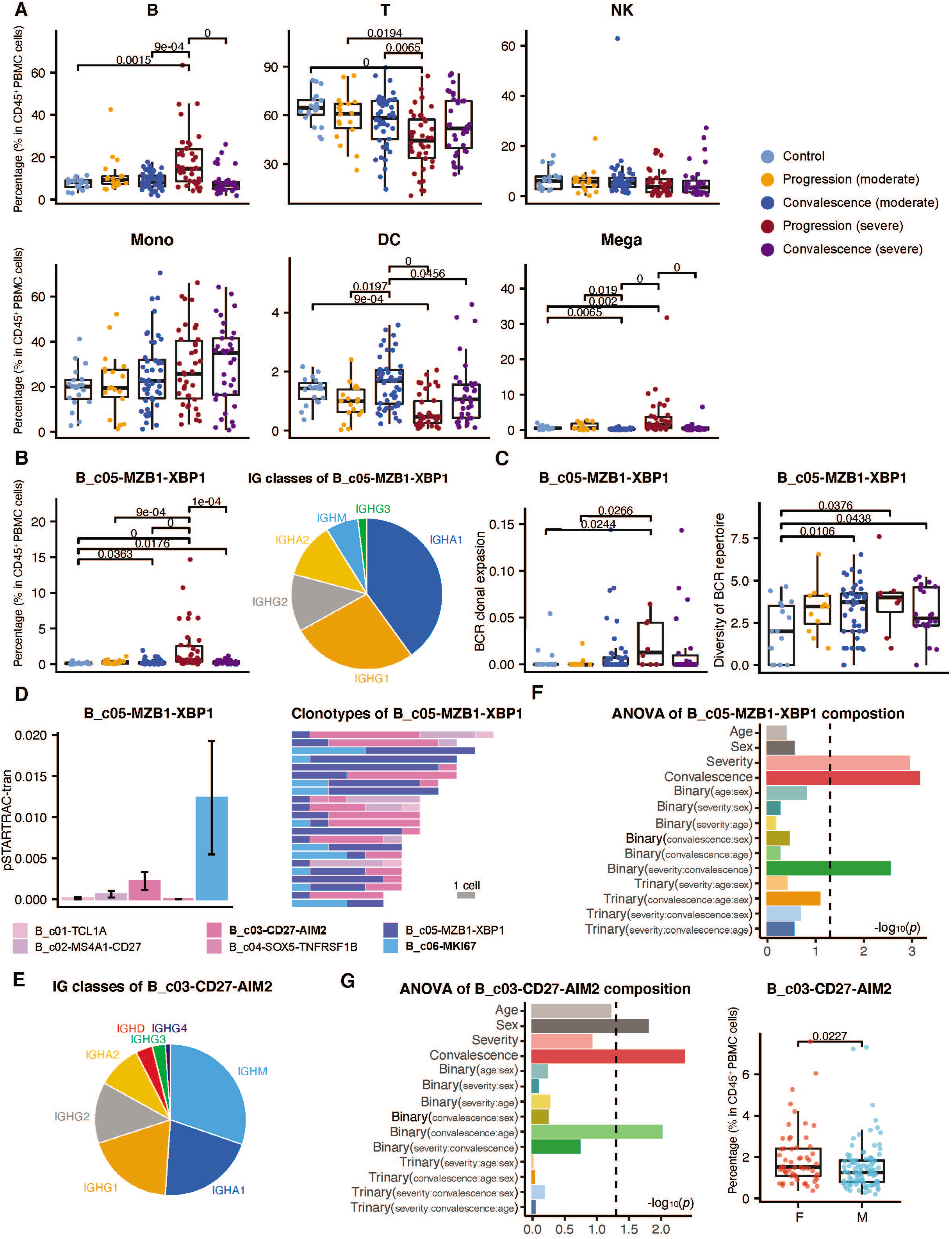
Dynamic changes of B cell composition across disease conditions. (A) Differences in immune cell composition across disease conditions for PBMC. Conditions are shown in different colors. Each boxplot represents one cell cluster. All differences with adjusted *P*-value < 0.05 are indicated; two-sided unpaired Wilcoxon was used for analysis. (B) Changes of XBP1+ plasma cells proportion across disease conditions. Composition of XBP1+ plasma cell BCRH cgene. All differences with *P*-value < 0.05 are indicated; twosided unpaired Wilcoxon was used for analysis. (C) Differences of XBP1+ plasma cells clonal expansion and BCR diversity across disease conditions. BCR clonal expansion level is calculated by STARTRAC-expa. Shannon’s entropy reveals the diversity of BCR repertoire. All differences with *P*-value < 0.05 are indicated; two-sided unpaired Wilcoxon was used for analysis. (D) Transition between XBP1+ plasma cells and other B cell sub clusters (left). Clonotypes of clones contain XBP1+ plasma cells (right); only shows clones with more than 5 cells. (E) Composition of B_c03-CD27-AIM2 memory cells BCRH cgene. (F) ANOVA of XBP1+ plasma cells proportion. (G) ANOVA of B_c03-CD27-AIM2 memory cells proportion (left) and differences of B_c03-CD27-AIM2 memory cells proportion between male and female (right). Two-sided unpaired Wilcoxon test. See also Figure S2.

### Elevated plasma B cells in COVID-19 patients

As single-cell dissection can provide the finest resolution to investigate the compositional changes among different COVID-19 patient groups, we then examined the heterogeneity of sub-clusters within each major immune cell type. For B cells, XBP1+ plasma cells (B_c05-MZB1-XBP1) showed the most significant compositional increases in PBMCs. For some severe COVID-19 patients, the percentage of plasma cells could even reach 15% of CD45+ cells in PBMCs, but the levels in other COVID-19 patients and healthy controls were less than 3% (Figure 2B). These cells highly expressed the genes encoding the constant regions of IgA1, IgA2, IgG1 or IgG2 (Figure 2B), indicating their functions to secrete antigen-specific antibodies to combat viral infection. This observation is consistent with the recent report that the serum of severe COVID-19 patients had high titers of SARS-CoV-2-specific antibodies (Tan et al., 2020b; Zhang et al., 2020c).

The increased plasma B cells in peripheral blood appeared to be derived from active proliferation of plasmablasts and transitions from memory B cells based on the paired BCR sequencing analyses. Both the extent of BCR clonal expansion and the diversity of the total BCR repertoire of these cells were significantly increased in severe COVID-19 patients (Figure 2C). Plasmablast cells (B_c06_MKI67), characterized by high expression of MKI67 and thus indicating a proliferative state, were elevated in the peripheral blood of severe COVID-19 patients (Figure S2A) and shared the most clonotypes with plasma cells (Figure 2D). The memory B cell cluster expressing high levels of *CD27, CD80, AIM2, GRIP2,* and *COCH* (B_c03-CD27-AIM2) was the second major source of plasma B cells in the peripheral blood, which shared a large proportion of clonotypes with plasma cells and plasmablasts (Figure 2D). Distinct from plasma cells and plasmablasts which were mainly composed of IgAs and IgGs, B_c03-CD27-AIM2 had a higher proportion of IgMs (Figure 2E), indicating a precursor state.

We applied analysis of variance (ANOVA) to dissect the associations of compositional changes of plasma B cells with disease severity, stage (progression or convalescence), age, sex, or the interactions of these factors. We found that plasma B cells in blood were specifically associated with the disease severity of COVID-19, and then disease stage, but had no associations with age or sex observed (Figure 2F) (Takahashi et al., 2020). In fact, for the mild/moderate disease, convalescent patients harbored higher levels of plasma B cells than those in the disease progression stage. By contrast, the plasma B cell levels in convalescent patients who recovered from severe disease were significantly lower than those in the disease progression stage (Figure 2B). Interestingly, the precursors of plasma B cells, *i.e.,* cells of B_c03-CD27-AIM2, appeared to be associated with sex differences (Figure 2G). In females, the percentage of B_c03-CD27-AIM2 cells was significantly higher than that of males (Figure 2G). Almost all B cell clusters were associated with disease stages, implying the importance of humoral immune response changes between disease progression and convalescence (Figure S2B and Table S3).

In summary, plasma B cells appeared to be significantly elevated in the peripheral blood regarding either the composition, proliferation, or developmental transition from memory B cells, and were more associated with disease severity. While their precursor cells were also elevated, they were more prone to be influenced by sex differences, providing a plausible explanation for the epidemiological observations on sex differences of COVID-19. Taken together with the observation that plasma B cells were more enriched in BALF (Figure 1F), these observations may suggest that humoral immune responses were actively initiated to combat SARS-CoV-2 infection and contributed to disease severity.

### Elevated proliferative T cells in COVID-19 patients

Similar to plasma B cells, proliferative CD8+ and CD4+ T cell clusters also showed an enrichment in BALF (Figure 1F) and elevation in PBMCs of COVID-19 patients albeit with a decrease of total T cells (Figures 2A and 3A) (Liao et al., 2020). A total of three proliferative CD8+ T cell clusters identified in this study, including T_CD8_c10-MKI67-GZMK, T_CD8_c11-MKI67-FOS, and T_CD8_c12-MKI67-TYROBP, were all increased in COVID-19 patients but with different characteristics. T_CD8_c10-MKI67-GZMK, a proliferative effector memory CD8+ T cell group characterized by high expression of *STMN1, HMGB2, MKI67,* and *GZMK,* was increased in the convalescent stage of severe COVID-19 patients (Figure 3B). Similarly, T_CD8_c11-MKI67-FOS also highly expressed *STMN1, HMGB2*, and *MKI67,* but exhibited low levels of *GZMK* instead and high levels of *FOS.* This cluster was increased in the disease progression stage of severe patients but not in convalescence (Figure 3B). T_CD8_c12-MKI67-TYROBP was characterized by high expression of *STMN1, HMGB2, MKI67,* and a NK cell marker gene *TYROBP.* This cluster was specifically increased in mild/moderate patients during the disease progression stage but deceased in the convalescence to a normal level as in healthy controls (Figure 3B). These observations were consistent with the activation of T cell responses in the peripheral blood of COVID-19 patients as previously reported using flow cytometry or CyTOF techniques (Mathew et al., 2020a; Sekine et al., 2020). However, the variations of proliferative CD8+ T cell clusters in different severity and stages have not been noticed before, which may indicate the complexity of T cell responses induced by SARS-CoV-2 infection in different patients. Moreover, in contrast to plasma B cells that accounted for 6.39% of total B cells, each proliferative CD8+ T cell cluster accounted for a much smaller proportion of the total CD8+ T cells (<1.63%) (Guo et al., 2020a; Liao et al., 2020; Mathew et al., 2020a).

**Figure 3.**
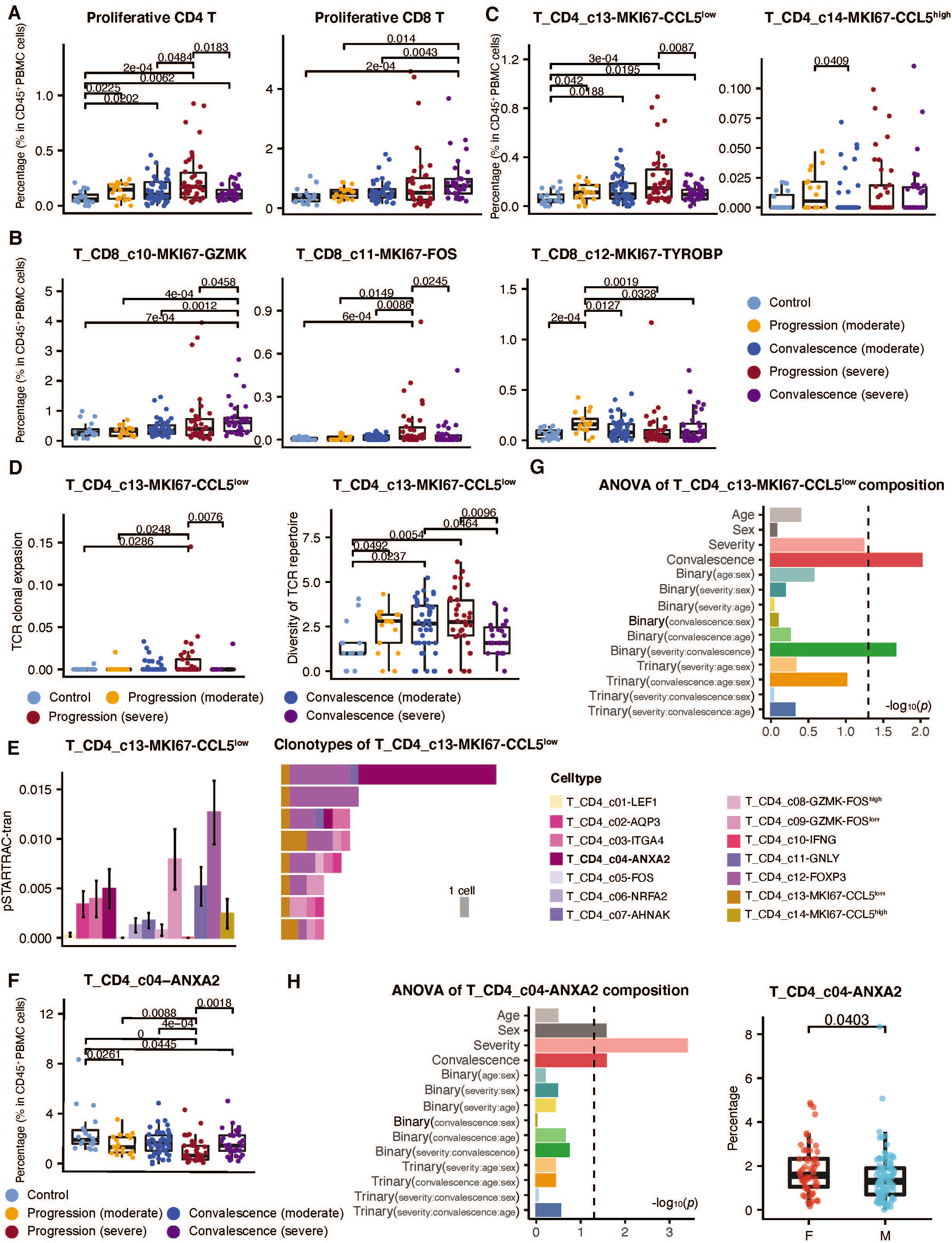
Differences in T cell composition across disease conditions. (A) Changes of proliferating CD4 and CD8 T cells across disease conditions for PBMC. Conditions are shown in different colors. All differences with adjusted *P*-value < 0.05 are indicated; two-sided unpaired Wilcoxon was used for analysis. (B) Differences of three CD8 proliferating T cell sub clusters proportion across disease conditions. All differences with *P*-value < 0.05 are indicated; two-sided unpaired Wilcoxon was used for analysis. (C) Differences of two CD4 proliferating T cell sub clusters proportion across disease conditions. All differences with *P*-value < 0.05 are indicated; two-sided unpaired Wilcoxon was used for analysis. (D) Differences of T_CD4_c13-MKI67-CCL5^low^ proliferating cells clonal expansion and TCR diversity across disease conditions. TCR clonal expansion level is calculated by STARTRAC-expa. Shannon’s entropy reveals the diversity of BCR repertoire. All differences with *P*-value < 0.05 are indicated; two-sided unpaired Wilcoxon was used for analysis. (E) Transition between T_CD4_c13-MKI67-CCL5^low^ proliferating cells and other CD4 cell sub clusters (left) and clonotypes of clones contain T_CD4_c13-MKI67-CCL5^low^ proliferating cells (right); only shows clones with more than 5 cells. (F) Differences of T_c04_CD4-ANXA2 T cell proportion across disease conditions. All differences with *P*-value < 0.05 are indicated; two-sided unpaired Wilcoxon was used for analysis. (G) ANOVA of T_CD4_c13-MKI67-CCL5^low^ proliferating cells proportion. (H) ANOVA of T_c04_CD4-ANXA2 T cell proportion (left) and differences of T_c04_CD4-ANXA2 T cell proportion between male and female (right). Two-sided unpaired Wilcoxon test. See also Figure S2.

Two proliferative CD4+ T cell clusters were also identified, with T_CD4_c13-MKI67-CCL5^low^characterized by high expression of *SELL* and low *CCL5* and T_CD4_c14-MKI67-CCL5^high^ characterized by low *SELL* and high *CCL5*. The counts of T_CD4_c14-MKI67-CCL5^high^ in PBMCs did not show significant differences among different COVID-19 patients. By contrast, the T_CD4_c13-MKI67-CCL5^low^ counts were elevated in COVID-19 patients, particularly in severe patients during the disease progression stage (Figure 3C). Similar to plasma B cells, the diversity and clonality of this cluster were both increased in severe patients with disease progression (Figure 3D), indicating an expanded TCR repertoire and developmental transitions from other clusters. Unlike plasma B cells whose source cluster B_c03-CD27-AIM2 was increased in peripheral blood (Figure S2C), the major source cluster of proliferative CD4+ T cells T_c04_CD4-ANXA2 was decreased in COVID-19 patients, particularly in severe patients during the disease progression stage (Figures 3E and 3F). This may partially explain the dichotomous and incomplete adaptive immunity previously observed in COVID-19 patients (Catanzaro et al., 2020). ANOVA analyses revealed that different from T_CD4_c13-MKI67-CCL5^low^ (Figure 3G), the percentage of T_CD4_c04-ANXA2 was associated with disease severity, progression/convalescence, and sex (Figure 3H). In particular, female patients generally had higher levels of T_CD4_c04-ANXA2 than males (Figure 3H), indicating the sex differences of T cell responses to SARS-CoV-2 infection (Takahashi et al., 2020).

In contrast to proliferative T cells that were elevated in PBMCs, other T cell clusters showed decrease in COVID-19 patients albeit with varied magnitudes, consistent with the lymphopenia that has been frequently observed in COVID-19 patients (Giamarellos-Bourboulis et al., 2020; Kuri-Cervantes et al., 2020; Yu et al., 2020). The most significantly decreased T cell clusters included γδT cells (T_c14_gdT-TRDV2), MAIT cells (T_CD8_c09-SLC4A10), a CD8+ T cell cluster highly expressing *TYROBP, KLRF1, CD247* and *IL2RB* (T_CD8_c08-IL2RB), and three CD4+ T cell clusters showing effector memory characteristics (T_CD4_c09-GZMK-FOS^low^, T_CD4_c11-GNLY, and T_CD4_c04-ANXA2). ANOVA analysis suggested that these clusters were mainly associated with disease severity rather than age or sex (Figures S2D and S2E and Table S3), implicating their critical roles in the disease progression of COVID-19. In particular, decreases of γδT cells and MAIT cells in the peripheral blood of COVID-19 patients (Figures S2F and Figure S2G) have been supported by flow cytometry-based analyses, suggesting their potent antimicrobial functions (Jouan et al., 2020).

While the decrease of γδT cells, MAIT cells, and effector memory T cells abovementioned were primarily associated with disease severity, the decreases of naive and central memory T cells were associated with the age but not sex difference of patients (Figure S2D and S2E and Table S3). Such clusters included the naive CD8+ cluster T_CD8_c01-LEF1, the CD8+ central memory cluster T_CD8_c02-GPR183, the naive CD4+ cluster T_CD4_c01-LEF1, and two CD69+ CD4+ clusters T_CD4_c06-NR4A2 and T_CD4_c05-FOS.

In summary, our scRNA-seq study recapitulated the lymphopenia in COVID-19 patients frequently observed in previous studies (Dhama et al., 2020; Giamarellos-Bourboulis et al., 2020; Kuri-Cervantes et al., 2020; Tan et al., 2020a; Yan et al., 2020; Yu et al., 2020). We further confirmed the activation of both CD4+ and CD8+ T cell responses in PBMCs recently found by flow cytometry-based immune profiling (Mathew et al., 2020a; Sekine et al., 2020). With the high resolution provided by scRNA-seq, we revealed the existence of distinct proliferative T cell clusters for both CD4+ and CD8+ T cells in COVID-19 patients, and implicated their different roles in patients of different groups and stages. Our large cohort also enabled us to dissect the impact of age and sex on the immune responses of COVID-19 patients. We found that, rather than associated with T cell proliferation, age and sex are more likely associated with the abundance of naive/central memory T cells and the precursor cells of proliferative T cells, respectively, highlighting the complexity of human T cell responses to SARS-CoV-2 infection.

### TCR/BCR usage patterns by COVID-19 patients

Our scRNA-seq data also coupled with TCR/BCR repertoire sequencing and thus provided a rich resource to investigate the TCR/BCR usage of COVID-19 patients, which is instructive for the development of anti-SARS-CoV-2 therapeutics and vaccines. We first examined whether identical TCRs or BCRs could be identified across COVID-19 patients. We found that only a few TCRs or BCRs were shared between two patients, and no identical TCRs or BCRs were shared beyond three patients. No TCRs or BCRs had identical amino acid sequences in more than three patients for the complementarity determining regions 3 (CDR3s) of β chains of TCRs or heavy chains of BCRs. We further examined whether the amino acid sequences of CDR3s of published SARS-CoV-2-reacting antibodies could be identified in the BCR repertoire of this cohort. We found that only one non-clonal BCR had identical CDR3 in its heavy chain with a comprehensive compendium containing 1,505 SARS-CoV-2-specific antibodies (Cao et al., 2020). Such scarcity of common TCRs or BCRs was in contrast with previous studies on severe patients recovered from enterovirus A71 infection and influenza vaccination (Chen et al., 2017; Jiang et al., 2013), suggesting that SARS-CoV-2 infection might not impose dramatic selective pressure on the somatic evolution of TCRs and BCRs.

Although no identical BCRs were found, we noticed that the BCR repertoire of COVID-19 patients had biased VDJ usage compared with that of healthy controls. We trained a random forest classifier with the VDJ usage frequencies to discriminate COVID-19 patients with mild/moderate or severe symptoms from healthy controls and found that the classification accuracy measured by the values of area under curve (AUC) could reach as high as 0.85. The most important VDJ combinations selected by the random forest classifiers also had significant overlaps with those of experimentally verified SARS-CoV-2-reacting antibodies (Figure S2H). Among the top 20 VDJ combinations important to discriminate severe COVID-19 patients from healthy controls selected by random forests, 14 had identical VDJ usage with experimentally verified neutralizing antibodies. Of note, the VDJ usage of the currently known SARS-CoV-2-neutralizing antibodies was biased towards IGHV3 and IGHV1. In particular, more than 40 neutralizing antibodies used IGHV3-53. Such observations are important to the development of effective diagnostics to trace human infection history and the further refinement of the current neutralizing antibodies.

The diversity of TCR or BCR repertoires of various T and B clusters might also be influenced by age, sex, COVID-19 severity, and disease stages. While age was mainly associated with the abundance of only naive and central memory T cells in PBMC, ANOVA analysis revealed that age might influence the decrease of TCR diversity in a wider range of T cells, including naive, central memory, and diverse effector memory T cells (Figure S2I-J). By contrast, sex differences were mainly associated with the BCR diversity of naive and memory B cells (B_c01-TCL1A, B_c02-MS4A1-CD27, and B_c04-SOX5-TNFRSF1 B) and the TCR diversity of a subset of effector memory CD4+ T cells (T_CD4_c08-GZMK-FOS^high^) (Figure S2I-K). After correcting the effects of age and sex, the decrease of diversity in MAIT cells, naive B and CD8+ and CD4+ T cells, effector memory CD8+ T cells (T_CD8_c03-GZMK and T_CD8_c04-COTL1), and a few CD69+ CD4+ T cell clusters (T_CD4_c03-ITGA4, T_CD4_c04-ANXA2) remained independently associated with COVID-19 severity (Figure S2I-K), highlighting the importance of these cells in COVID-19. Importantly, the TCR diversity of one proliferative CD8+ T cell cluster, *i.e.,* T_CD8_c11-MKI67-FOS, was associated with the triad interaction by disease severity, age, and sex (Figure S2J), indicating the impacts of age and sex on disease severity. Similarly, the clonal expansion of a central memory CD4+ T cell cluster highly expressing *AQP3* (T_CD4_c02-AQP3) was also associated with the triad interaction by disease severity, age, and sex (Table S3), indicating that age and sex might impact the COVID-19 disease via multiple mechanisms.

Taken together, our data suggested that SARS-CoV-2 might not impose dramatic selective pressure on the somatic evolution of TCRs or BCRs for COVID-19 patients, thus resulting in few identical TCRs and BCRs across patients. However, preferential VDJ usage were identified, which highly overlapped with the sequences of some known SARS-CoV-2-neutralizing antibodies. The diversity of TCR and BCR repertoires of various T and B clusters might be shaped by age, sex, disease severity, and stages together, although the influences of these factors were heterogeneous on different cell types. In particular, triad interactions among age, sex and disease severity were indicated for specific CD8+ and CD4+ clusters, underscoring the complex T cell responses of COVID-19 patients and providing important clues for future studies.

### SARS-CoV-2 detected in multiple epithelial and immune cell types with interferon response phenotypes

The enrichment of plasma B and proliferative T cells in BALF and the elevation of these cells in PBMCs of COVID-19 patients highlighted the roles of these cells in combating SARS-CoV-2 infection. To explore potential interactions between these cells and SARS-CoV-2 infected cells, we examined the characteristics of cell types that harbored SARS-CoV-2 sequences in our dataset. From the BALF samples of severe COVID-19 patients in the disease progression stage, we identified viral RNAs of SARS-CoV-2 in three epithelial cell types including ciliated, secretory, and squamous epithelial cells and a diverse set of immune cells including neutrophils, macrophages, plasma B cells, T cells, and NK cells (Figure 4A). The cell identities of these SARS-CoV-2-positive cells were well confirmed by their corresponding molecular markers (Figure 4B), excluding the possibility of artefacts caused by doublets during scRNA-seq. Because ACE2 and TMPRSS2 have been recognized to play critical roles in mediating viral entry into the host cells for SARS-CoV-2, we examined their expression levels in these cells (Figure 4C) (Netea et al., 2020). We found that at least a subset of those epithelial cells expressed *ACE2* and *TMPRSS2*, consistent with the notion that SARS-CoV-2 employs ACE2 and TMPRSS2 to invade these cells. Interestingly, those immune cells, which did not express *ACE2* or *TMPRSS2,* harbored even more viral RNA sequences than the epithelial cells (Figure 4D). The high viral load reassured that the detection of SARS-CoV-2 RNAs in these immune cells was unlikely caused by experimental contamination. Consistently, an independent scRNA-seq study of COVID-19 patients also identified SARS-CoV-2 RNAs in neutrophils and macrophages from the respiratory samples of COVID-19 patients (Bost et al., 2020).

**Figure 4.**
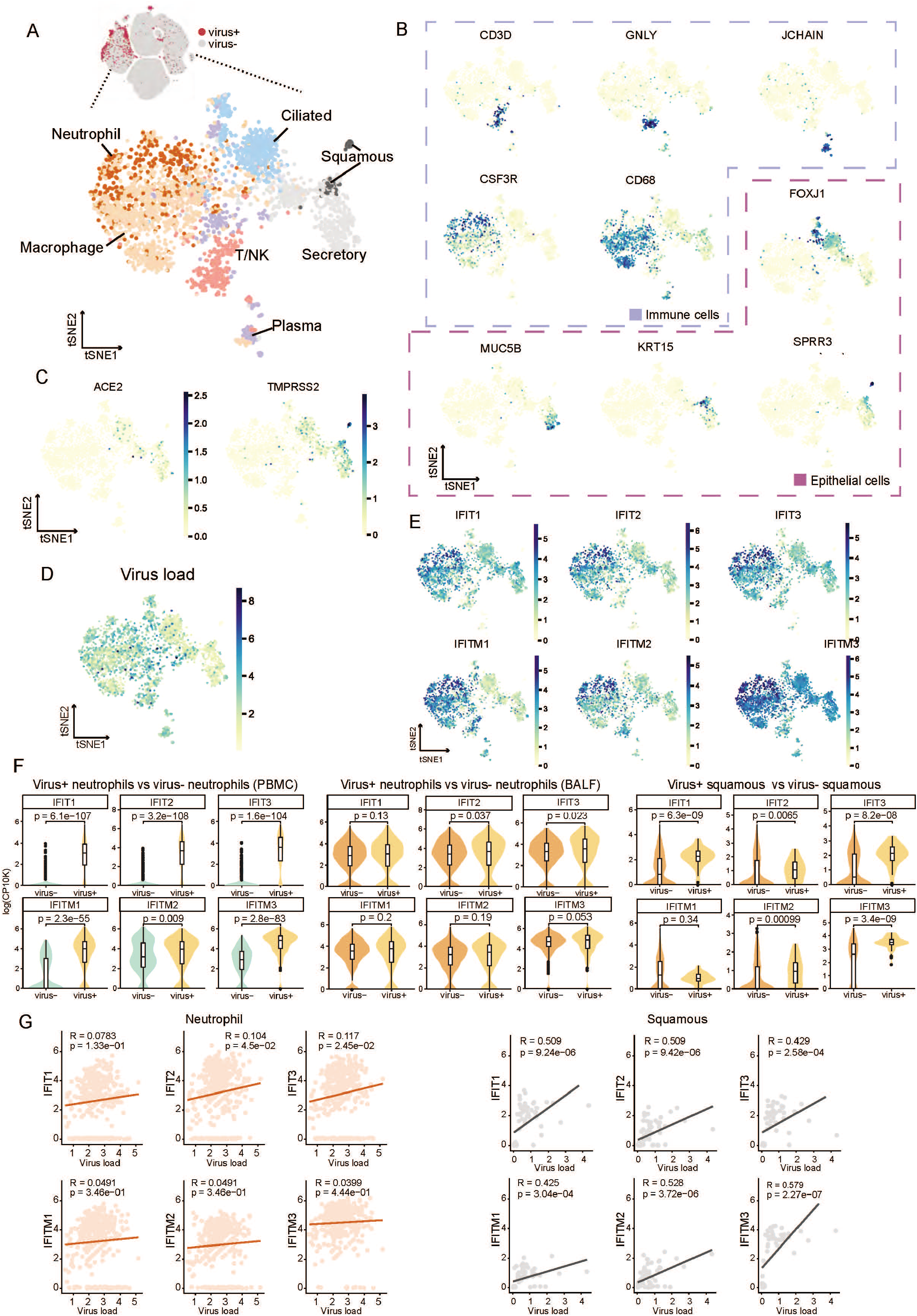
Landscape Of Cell Types Detected SARS-Cov-2 Sequences and Their Antiviral Response. (A) Uniform Manifold Approximation and Projection (UMAP) of all cells with SARS-CoV-2 genome UMI > 0 after quality control containing 3085 cells in total. (B) Characteristic markers we chose to identify each cell type. The purple box indicates immune cell types (top), and the red one indicates epithelial cell types (bottom). (C) UMAP showing expression level of known SARS-CoV-2 infected receptor ACE2 (left) and TMPRSS2 (right). Each dot denotes a single cell and colored by its expression level of the gene. (D) UMAP showing the viral load of each cell. The darker colors in the bar indicate a higher viral load in cells. (E) UMAP showing the activation of Interferon-stimulated genes (ISGs) in cells with viral detection. (F) Violin plots showing differential expression of ISGs between cells with viral detection (virus+) and cells without (virus-) in PBMC-derived neutrophils (left panel), BALF-derived neutrophils (middle panel) and squamous cells (right panel). The y axis represents the expression level of each gene. logCP10K, log-transformed counts per 10,000. Two-sided unpaired Wilcoxon test. (G) Scatter plots showing the correlation between viral load and ISGs in neutrophil (left panel) and squamous cells (right panel). The line in scatter plots represent the result of linear regression. Each point in the graph represents one single cell, colored by cell types. The x axis shows virus load in each cell while the y axis represents the expression level of one of the ISG genes. Correlation coefficient (R) and probability (*p*) are acquired using Pearson’s correlation. See also Figures S3 and Tables S4.

Since interferon-stimulated genes (ISGs) are typically activated in virus-infected cells (Schoggins and Rice, 2011), we next examined the expression of ISGs in these cells (Figure 4E and Table S4). Because *IFIT1/2/3* and *IFITM1/2/3* are frequently observed to increase after various viral infections (Zhang et al., 2016), the high expression of these genes in these immune cells, particularly neutrophils and macrophages, may indicate ISG activation in these cells. Compared with matched cell types in PBMC, almost all these *ISG* genes exhibited elevated expression in these virus+ immune cells (Figures 4F and S3B and Table S5). Compared with virus-negative immune cells of the same types in the BALF, SARS-CoV-2+ epithelial cells, including ciliated, secretory, and squamous cells, as well as those virus-positive neutrophils, exhibited higher levels of *ISG* expression (Figure 4F and Table S5). Positive correlations between the viral loads estimated by the abundance of viral RNAs and the *ISG* expression levels were observed for squamous epithelial cells but not ciliated or secretory epithelial cells (Figures 4G and S3C). For immune cells, virus-positive neutrophils exclusively demonstrated positive correlations between viral loads and *ISG* levels (Figure 4G), but this phenomenon did not exist in other immune cell types. These observations suggest that SARS-CoV-2 might be able to infect human cells beyond traditionally assumed respiratory epithelial cells and could induce interferon responses.

In BALF from mild/moderate COVID-19 patients, fewer cells were obtained and no SARS-CoV-2 RNAs were detected in cells from such samples, suggesting that the respiratory tract of mild/moderate patients might be more intact and the viral titer was lower than those from severe patients. Although type II alveolar (AT2) cells were reported vulnerable to SARS-CoV-2 infection (Hou et al., 2020), our study revealed few AT2 cells in the BALF and no detectable SARS-CoV-2 RNAs in AT2 cells, which is consistent with the previous finding that lower respiratory tract cells had lower potential to be infected by SARS-CoV-2 than those from nasal and upper respiratory tract (Hou et al., 2020; Sungnak et al., 2020).

### Distinct transcriptomic changes of ciliated, secretory, and squamous epithelial cells after SARS-CoV-2 infection

SARS-CoV-2 infection in different epithelial cells resulted in not only distinct interferon responses, but also significant transcriptomic changes. For squamous epithelial cells, SARS-COV-2+ cells exhibited elevated expression of a diverse set of genes such as *NT5E, CLCA4,* and *SULT2B1* (Figure 5A). These genes were enriched in pathways such as “response to virus”, “response to type I interferon” and “response to hypoxia”, consistent with viral infection and the subsequent respiratory distress, reflecting the host immune response via type I interferons (Figure 5B). By contrast, the numbers of genes with significant changes after SARS-CoV-2 infection for ciliated and secretory epithelial cells were much smaller than squamous cells, and few genes showed consistent changes in all the three epithelial cell types (Figure 5C).

**Figure 5.**
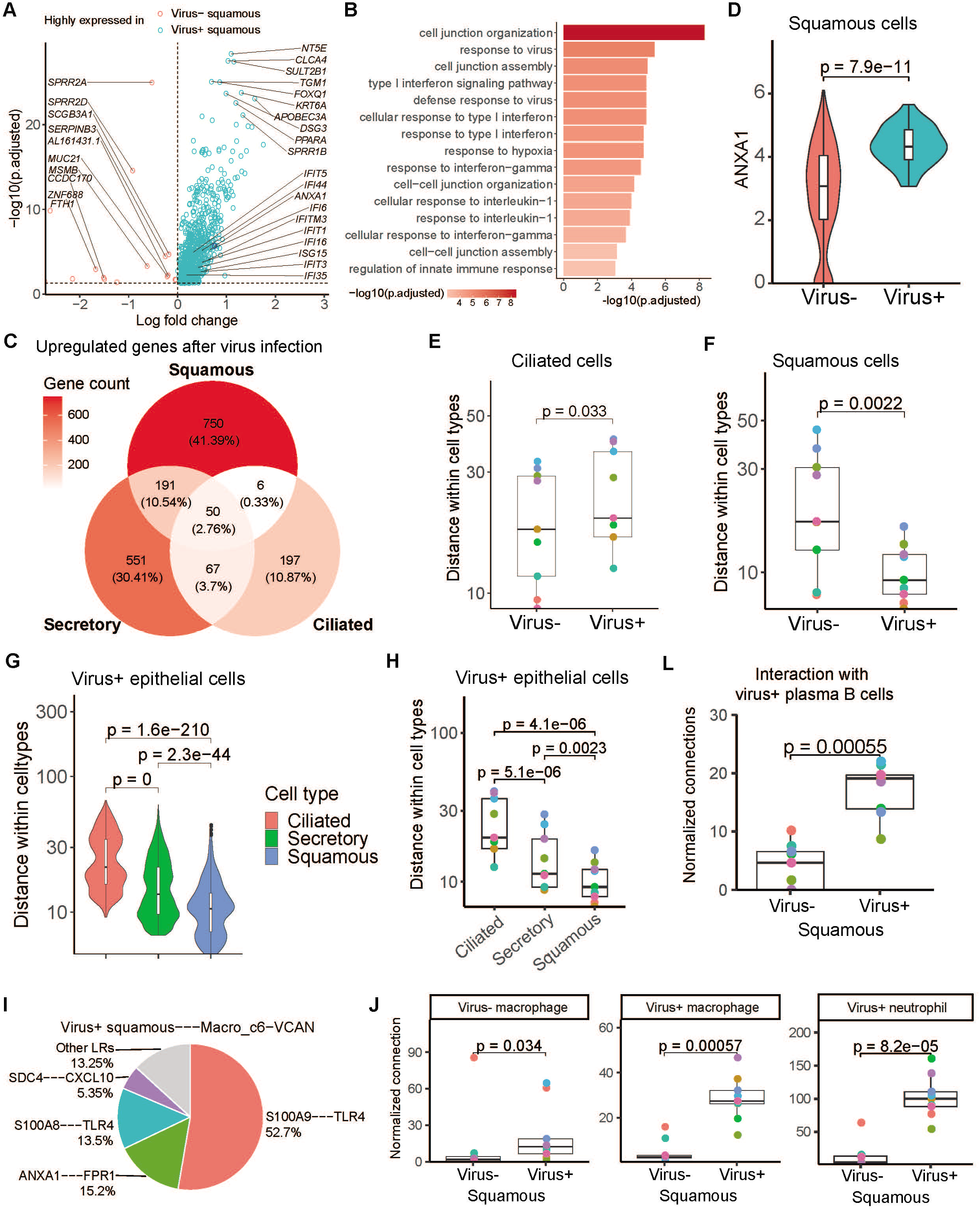
The Impact of Virus Infection on Expression and Cell-cell interaction of Epithelial Subtypes. (A) Volcano plot showing differentially expressed genes between squamous cells with or without viral detection. Adjusted *P*-value < 0.05, Two-sided unpaired Wilcoxon test. *ANXA1* is denoted in dark blue. (B) Enriched GO terms of genes highly expressed in virus+ squamous cells shown in (A). (C) Venn plot showing the intersection of genes upregulated in epithelial cells with viral detection. Each compartment is colored by the number of genes. (D) Violin plot showing the expression of *ANXA1* in squamous cells with or without viral detection. Two-sided unpaired Wilcoxon test. (E) Boxplot showing the pseudo space distance within ciliated cells. Each dot represents an individual patient. Two-sided paired Wilcoxon test. (F) Boxplot showing the pseudo space distance within squamous cells. Each dot represents an individual patient. Two-sided paired Wilcoxon test. (G) Violin plot showing the pseudo space distance within each type of epithelial cells in one example. Two-sided unpaired Wilcoxon test. (H) Boxplot showing the median of pseudo space distance within each type of epithelial cells of all the patients with BALF data. Each dot represents an individual patient. Two-sided unpaired Wilcoxon test. (L) Boxplot showing the normalized connection between squamous cells and virus-detected plasma B cells of all the patients with BALF data. Each dot represents an individual patient. Two-sided unpaired Wilcoxon test. (I) Pie chart showing the ligand-receptor contribution proportion between virus+ squamous and Macro_c6-VCAN in one example. Ligand-receptor pairs with contribution less than 0.05 were merged as ‘Other LRs’. (J) Boxplot showing the normalized connection between squamous cells and virusmacrophage (left), virus+ macrophage (middle) and virus+ neutrophils (left). Each dot represents an individual patient. Two-sided unpaired Wilcoxon test. See also Figures S4 and S5.

We next explored the impact of the above transcriptomic changes, especially on their interaction potentials with immune cells. Annexin A1 (*ANXA1*), up-regulated in virus+ squamous epithelial cells (Figure 5D), is known to regulate the functions of neutrophils in inflammation via its interactions with formyl peptide receptors (Sugimoto et al., 2016). This prompted us to investigate the cellular interaction changes of epithelial cells with each other and with immune cells after SARS-CoV-2 infection. Based on CSOmap that estimates cellcell interactions in three-dimensional space via ligand-receptor (LR) mediated cell selforganization and competition (Ren et al., 2020), we estimated the cellular interaction potentials in a computationally constructed pseudo-space and found that ciliated, secretory, and squamous epithelial cells exhibited distinct interaction potentials after SARS-CoV-2 infection.

Ciliated epithelial cells exhibited lower interaction potentials with themselves and other cells after SARS-CoV-2 infection, and thus would disperse in the outer compartment of the pseudo-space (Figures 5E, S4A and S4B), consistent with the pathological phenomenon of epithelial denudation of coronavirus infection in respiratory tract (Lee et al., 2003; Nicholls et al., 2003). By contrast, squamous epithelial cells significantly enhanced their interacting potentials with themselves after SARS-CoV-2 infection compared with those squamous cells with no viral detection (Figure S4C). Such changes were consistent across COVID-19 patients (Figure 5F). Comparison across ciliated, secretory, and squamous epithelial cells infected by SARS-CoV-2 also highlighted the dispersing tendency of ciliated cells and the interacting potentials among squamous cells themselves (Figures 5G and 5H).

Such interaction distinctions not only existed among epithelial cells, but also impacted their interactions with immune cells. Consistent with the dispersing nature of ciliated cells in the outer compartment of the pseudo-space, no significant interactions were observed between virus+ ciliated cells and immune cells. By contrast, virus+ secretory epithelial cells showed significant interactions with neutrophils and macrophages in mild/moderate COVID-19 patients via the SCGB3A1-MARCO axis (Figures S4D and S4E), but such interactions were subdued in severe COVID-19 patients due to the down-regulation of *MARCO* in neutrophils and macrophages (Figure S4F). In severe patients, virus+ squamous cells showed significant interactions with neutrophils and macrophages via the ANXA1-FPR1 and S100A9/A8-TLR4 axes (Figure 5I). Neutrophils and macrophages exhibiting high interacting potentials with virus+ squamous epithelial cells were also prone to be SARS-CoV-2 infected (Figure 5J). As ANXA1-FPR1 and S100A9/A8-TLR4 interactions have been reported to play critical roles in the recruitment of immune cells and inflammatory cascade under various conditions including sepsis and tumor (Gavins et al., 2012; Laouedj et al., 2017; Osei-Owusu et al., 2019; Vogl et al., 2007), they might also play important roles in the pathogenesis of SARS-CoV-2 infection. In contrast to innate immune cells such as neutrophils and macrophages, T and B cells did not show significant interactions with any of the three types of virus+ epithelial cells (Figure S4G), implying a compromised adaptive immune response. It is noteworthy that plasma B cells in BALF also tended to be SARS-CoV-2-positive and displayed close interactions with virus+ neutrophils and squamous epithelial cells via the S100A9/A8-TLR4 axes (Figure 5L).

We then investigated the cell types expressing *ANXA1, FPR1, S100A9, S100A8,* and *TLR4* in both BALF and PBMC across COVID-19 patients to evaluate the possible inflammatory cascade mediated by these LR pairs. It was evident that *ANXA1* was highly expressed in a wide range of immune cells except B cells and naive T cells (Figures S5A and S5B) and its receptor *FPR1* was highly expressed in neutrophils, macrophages, and monocytes (Figures S5A and S5B). Interestingly, for most immune cell clusters in BALF, the expression levels of *ANXA1* and *FPR1* were down-regulated in severe COVID-19 patients compared with those of mild/moderate COVID-19 patients (Figure S5A). But in PBMCs, except for MAIT cells (T_CD8_c09-SLC4A10) and γδT cells (T_gdT_c14-TRDV2), *ANXA1* and *FPR1* were significantly up-regulated in many cell types in severe COVID-19 patients compared with those of mild/moderate COVID-19 patients (Figure S5B). *S100A9* and *S100A8* were highly expressed in neutrophils, macrophages, and monocytes in COVID-19 patients with mild/moderate symptoms and had no expression in T, B, NK, or dendritic cells (Figures S4H and S5B). However, for severe COVID-19 patients in the disease progression stage, *S100A9* and *S100A8* were significantly up-regulated in almost all cell clusters for both BALF and PBMCs (Figures S4H and S5B). In particular, T, B, NK, and dendritic cells had no or minimal levels of *S100A9* and *S100A8* expression in mild/moderate COVID-19 patients (Figures S4H and S5B). By contrast, in severe COVID-19 patients, the levels of *S100A9* and *S100A8* were significantly up-regulated in T, B, NK, and dendritic cells (Figures S4H and S5B), indicating a systemic inflammatory response. *TLR4* did not exhibit significant differences in PBMCs between severe and mild/moderate COVID-19 patients but was significantly down-regulated in certain BALF monocyte and macrophage subsets (Figure S5B).

In summary, our data indicated that SARS-CoV-2 infection in different types of epithelial cells might trigger different transcriptomic changes and thus could modulate their interactions with themselves and with immune cells. In particular, squamous epithelial cells could up-regulate *ANXA1* and *S100A8/A9* after SARS-CoV-2 infection, enhancing their interactions with neutrophils and macrophages via the axes of ANXA1-FPR1 and S100A8/A9-TLR4. The systemic up-regulation of *ANXA1, FPR1,* and *S100A8/A9* in immune cells from peripheral blood may indicate, at least partially, the molecular mechanism of aberrant inflammation in severe COVID-19 patients. This hypothesis is supported by a preliminary finding that small molecules targeting S100A8/A9 could inhibit SARS-CoV-2-induced aberrant inflammation in mice (Guo et al., 2020b). Thus, S100A8/A9 should be further evaluated as therapeutic targets. Compared to innate immune cells, adoptive immune cells including T and B cells did not show significant interactions with SARS-CoV-2-positive epithelial cells in BALF by computational simulation, consistent with previous findings (Chua et al., 2020; Wauters et al., 2020). These might suggest a compromised adaptive immune response in severe patients. Furthermore, the megakaryocytes in PBMCs, followed by monocytes, exhibited higher interaction potentials with epithelial and immune cells in BALF than adaptive immune cells (Figure S4I), suggesting the critical roles of these cells in the pathogenesis of COVID-19.

### Megakaryocytes and monocyte subsets as critical peripheral sources of cytokine storms

With our large scale scRNA-seq dataset, we next sought to investigate whether any crucial cell subtypes in peripheral blood contribute to the bulk of inflammatory cytokine production. We first defined a cytokine score and inflammatory score for each cell based on the expressions of the collected cytokine genes and reported inflammatory response genes (Liberzon et al., 2015) (Table S6), respectively, and used these two scores as indicators to evaluate the levels of inflammatory cytokine storm for each cell. We found apparent elevated expression of cytokine and inflammatory genes in patients, especially at the severe progression stage (Figures 6A and S6A), indicating the existence of inflammatory cytokine storm after SARS-CoV-2 infection. Seven cell subtypes, including three subtypes of monocytes (Mono_c1-CD14-CCL3, Mono_c2-CD14-HLA-DPB1 and Mono_c3-CD14-VCAN), three subtypes of T cells (T_CD4_c08-GZMK-FOS^high^, T_CD8_c06-TNF and T_CD8_c09-SLC4A10) and one subtype of megakaryocytes was detected with significantly higher cytokine and inflammatory scores (Figure. S9B, Table S2, *P* < 0.0001), indicating that these cells might be major sources of inflammatory storm. Interestingly, megakaryocytes, which have not been reported in the inflammatory response in COVID-19 patients, may affect the functions of platelets at the disease stage, in consistent to a previous study (Manne et al., 2020).

**Figure 6.**
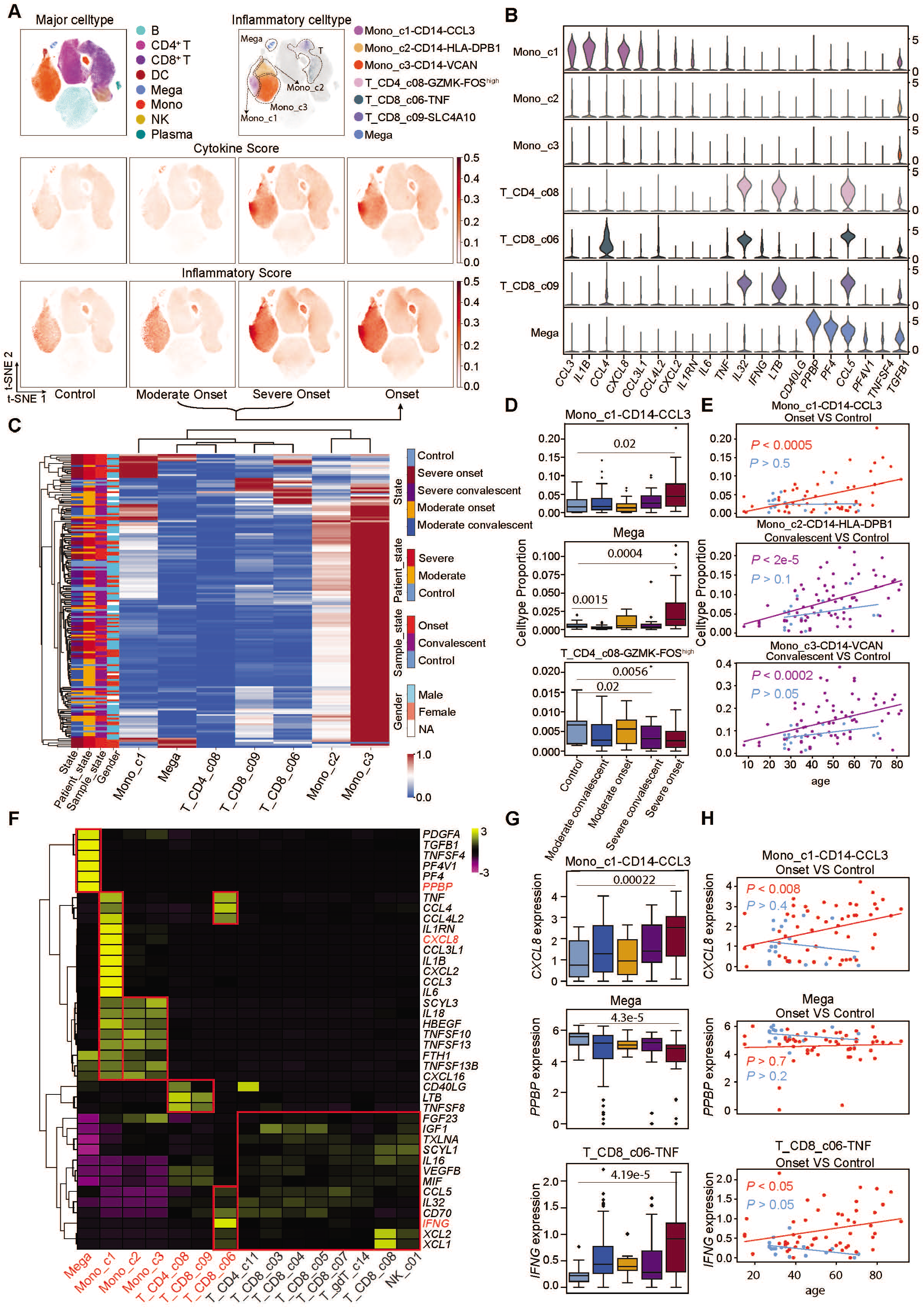
Mono_c1-CD14-CCL3 and megakaryocytes in peripheral blood appear as dominant source for inflammatory cytokine storm. (A) t-SNE plots of PBMC cells colored by major cell types (top left panel), inflammatory cell types (top right panel), cytokine score (middle panel) and inflammatory score (bottom panel). (B) Violin plots of selected cytokine genes for seven hyper-inflammatory cell subtypes. (C) Heatmap of an unsupervised clustering of cell proportion of seven hyper-inflammatory cell subtypes in all samples analyzed. (D) Box plots of the cell proportion of Mono_c1-CD14-CCL3, Mega and T_CD4_c08-GZMK-FOS^high^ clusters from healthy controls (n=20), moderate convalescent (n=48), moderate onset (n=18), severe convalescent (n=35) and severe onset (n=38) patients. Two-sided Wilcoxon rank-sum test. (E) Ordinary least squares model of age to cell proportion of Mono_c1-CD14-CCL3, Mono_c2-CD14-HLA-DPB1 and Mono_c3-CD14-VCAN clusters from healthy controls (n=20), convalescent (n=83) and onset (n=56) patients. P value was assessed with F-statistic for ordinary least squares model. (F) Heatmap of cytokines genes’ expression among seven hyper-inflammatory cell subtypes. Seven hyper-inflammatory cell subtypes are colored in red and others are colored in grey. (G) Box plots of the cytokines’ expression of Mono_c1-CD14-CCL3, Mega and T_CD8_c06-TNF clusters from healthy controls (n=20), moderate convalescent (n=48), moderate onset (n=18), severe convalescent (n=35) and severe onset (n=38) patients. Two-sided Wilcoxon rank-sum test. (H) Ordinary least squares model of age to cytokines’ expression of Mono_c1-CD14-CCL3, Mega and T_CD8_c06-TNF clusters from healthy controls (n=20), convalescent (n=48+35) and onset (n=18+38) patients. P value was assessed with F-statistic for ordinary least squares model. In (D) and (G), the box represents the second, third quartiles and median, whiskers each extend 1.5 times the interquartile range; dots represent outliers. In panel (B), (C) and (F), Mono_c1, Mono_c2, Mono_c3, T_CD4_c08, T_CD8_c09, T_CD8_c06 and Mega correspond to Mono_c1-CD14-CCL3, Mono_c2-CD14-HLA-DPB1, Mono_c3-CD14-VCAN, T_CD4_c08-GZMK-FOS^high^, T_CD8_c09-SLC4A10, T_CD8_c06-TNF and Mega, respectively. In panel (F), T_CD4_c11, T_CD8_c03, T_CD8_c04, T_CD8_c05, T_CD8_c07, T_gdT_c14, T_CD8_c08, NK_c01 correspond to clusters of T_CD4_c11-GNLY, T_CD8_c03-GZMK, T_CD8_c04-COTL1, T_CD8_c05-ZNF683, T_CD8_c07-TYROBP, T_gdT_c14-TRDV2, T_CD8_c08-IL2RB and NK_c01-FCGR3A, respectively. DC, dendritic cells. Mega, megakaryocytes. Mono, monocytes.

Each of the hyper-inflammatory subtypes highly expressed several cytokine genes that are known to be involved in the inflammatory storm, such as *CCL3, IL1B, CXCL8, CCL4, CCL6, IL32, LTB* and *TGFB1,* but with different patterns (Figure 6B), suggesting divergent genomic signatures of these cells. We then investigated the proportion of each of the 7 cell subtypes in every patient and found that these hyper-inflammatory cell subtypes were in general slightly more frequent in patients at severe stage (Figure. S6C). When we clustered these cell subtypes with each individual patient based on the proportions of the hyper-inflammatory cell subtype in PBMCs, we found distinct enrichment of these cell subtypes in different groups of patients (Figure 6C). Mono_c1-CD14-CCL3, known be associated with tocilizumab-responding cytokine storm (Guo et al., 2020a), was highly enriched in a subpopulation of severe onset patients likely to be accompanied by inflammatory storm (Figures 6C and 6D). The proportion of Mono_c1-CD14-CCL3 subtype was also correlated with the age of the corresponding patients (Figure. 6E). The hyper-inflammatory megakaryocytes were enriched in another batch of severe onset patients, which could also be under excessive inflammatory response (Figure. 6C and 6D).

By contrast, Mono_c2-CD14-HLA-DPB1 and Mono_c3-CD14-VCAN subtypes were widely distributed in every disease stage, and the hyper-inflammatory T cells showed decreased proportions in patients at the severe onset stage such as T_CD4_c08-GZMK-FOS^high^ subtype (Figures 6C, 6D and S6B), although both of these two monocyte subtypes exhibited increased proportions in elder convalescent patients (Figure 6E). Taken together, these results suggest that Mono_c1-CD14-CCL3 and megakaryocytes were the major sources triggering cytokine inflammatory storm, with both elevated cell ratios and inflammatory scores in the severe onset patients. On the other hand, although the severity of COVID-19 is correlated with lymphopenia, partially reflected by reduced T cells in PBMC (Dhama et al., 2020; Tan et al., 2020a; Yan et al., 2020), certain T cell subtypes might actually contribute to the inflammatory storm by enhanced expressing of proinflammatory cytokines.

Next, we investigated the inflammatory signatures for each hyper-inflammatory cell subtype and found unique pro-inflammatory cytokine gene expressions in each cell subtype (Figure 6F), suggesting diverse mechanisms by which these cell subtypes may contribute to the cytokine storm. The hyper-inflammatory Mono_c1-CD14-CCL3 and megakaryocytes largely expressed more cytokines, suggesting central roles of the two cell types in driving the inflammatory storm. Specifically, Mono_c1-CD14-CCL3 highly expressed *CXCL8, TNF, IL1RN, IL1B,* and *CCL3*, which we also detected with significantly higher levels in serum from patients at the severe stage, especially those critically ill patients (Figures 6F and S6D). Although the inflammatory megakaryocytes highly expressed the cell type identity marker genes such as *PPBP (Zhang et al., 1997),* the expression level of these genes was significantly decreased in patients compared to healthy controls, indicating a loss of function of these cells after inflammatory activation (Figures 6F and 6G). Notably, the T_CD8_c06-TNF subtype specifically and highly expressed *IFNG,* a pro-inflammatory cytokine highly enriched in patients at the severe onset stage also confirmed by serum cytokine detection (Figures 6F, G and S6D). Moreover, pro-inflammatory cytokines *CXCL8* and *IFNG* showed significant age-dependent expressions in patients with disease progression, while no significance was observed in healthy controls (Figure 6H). *PPBP* showed no correlation with the age in either patients or healthy controls, suggesting that the loss of function of megakaryocytes might not be age-dependent (Figure 6H). To assess the dynamic changes of cytokines in COVID-19 patients with different periods, we compared them with healthy controls for these seven hyper-inflammatory subtypes, and found that *IFNG, IL6, CCL3, TNF, CXCL2, CXCL8, IL1RN,* etc, were highly expressed in cells of severe patients with disease progression (Figure S6E).

We also observed eight cell subtypes with significantly higher cytokine scores even though their inflammatory scores showed no difference to other cell clusters (Figure S6B, Table S7, *p* < 0.0001). These cell subtypes exhibited uniform and relatively low expressions of cytokine genes such as *IGF1, TXLNA, SCYL1, CCL5* and *IL16* (Figure 6F), likely not involved in the cytokine storm. No significant differences were observed at the serum level for these cytokines between the different groups of patients (Figure S6F). These genes specific for hyper-inflammatory cells may serve as signatures for the inflammatory storm and be helpful in deepening the understanding of COVID-19 pathogenesis.

### Interactions of hyper-inflammatory cell subtypes in lung and peripheral blood

The dysregulated cytokine responses associated with the inflammatory cytokine storm may cause immunopathological injury to the lung, and large amount of infiltrating inflammatory immune cells have been demonstrated in the pulmonary tissue of COVID-19 patient (Bhaskar et al., 2020; Cao, 2020; Sun et al., 2020). We analyzed the expressions of cytokines and inflammatory genes for each cell from the BALF samples, and compared the inflammatory and cytokine scores among all the cell subtypes captured in BALF. No enrichment of cytokine genes was observed from the epithelial cells, while subtypes of macrophages and monocytes had the highest cytokine and inflammatory scores in the severe onset samples (Figure 7A). Similar to our analysis on PBMCs, we identified five hyper-inflammatory cell subtypes, including Macro_c2-CCL3L1, the three subtypes of monocytes and the neutrophils (Figure 7B), suggesting that these cell subtypes might be the major sources driving inflammatory storm in the lung tissue. Neither CD4+ nor CD8+ T cells were detected with an elevated inflammatory score or the cytokine score in BALF samples, which was different from those in PBMCs. Each hyper-inflammatory subtype highly expressed specific cytokines; for example, Macro_c2-CCL3L1 specifically expressed *CCL8*, *CXCL10/11,* and *IL6*. Mono_c1-CD14-CCL3, as one of the most notable proinflammatory cell types in both peripheral blood and BALF, uniquely expressed high expression levels of *IL1B, CCL20, CXCL2, CXCL3, CCL3, CCL4, HBEGF* and *TNF*. The neutrophils also showed many uniquely expressed cytokines including *TNFSF13B, CXCL8, FTH1, CXCL16* (Figure 7C).

**Figure 7.**
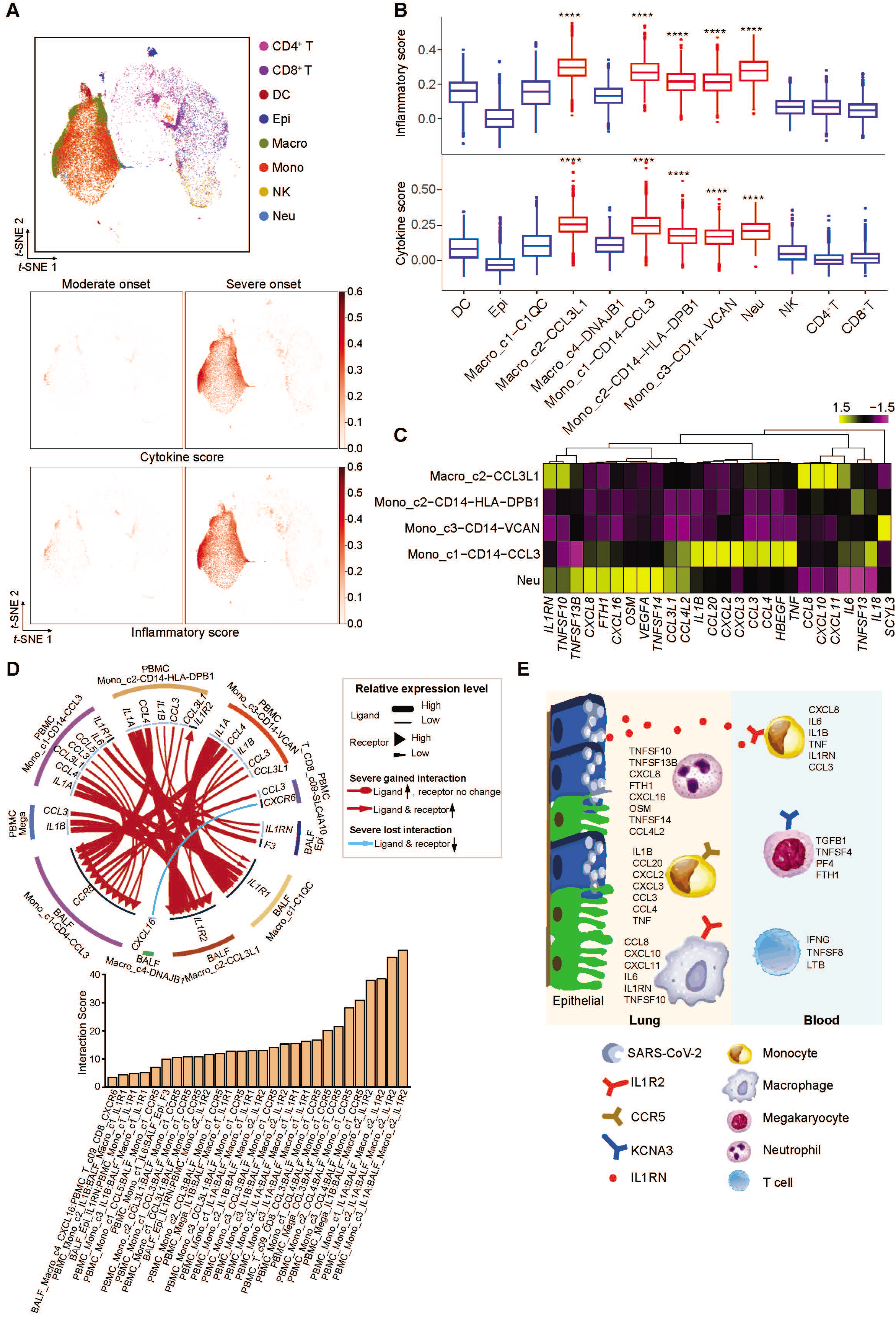
The interactions of hyper-inflammatory cell subtypes in lung and peripheral blood. (A) t-SNE plots of BALF cells colored by major cell types (top panel), cytokine score (middle panel) and inflammatory score (bottom panel). (B) Boxplots of the inflammatory score (top panel) and cytokine score (bottom panel) of cell subtypes. Significance was evaluated with Wilcoxon rank-sum test. **** P < 0.0001. (C) Heatmap of an unsupervised clustering of cytokine genes’ expression among five hyper-inflammatory cell subtypes. (D) Circos plot showing the prioritized interactions mediated by ligand-receptor pairs between inflammation-related cell types from BALF and PBMC, respectively. The outer ring displays color coded cell types and the inner ring represents the involved ligand-receptor interacting pairs. The line width and arrow width are proportional to the log fold change between severe onset and moderate onset patient groups in ligand and receptor, respectively. Colors and types of lines are used to indicate different types of interactions as shown in the legend. The bar plot at bottom indicates the interaction score for each interaction which serves to measure the interaction strength. (E) Summary illustration depicting the potential cytokine/receptor interactions of hyper-inflammatory cell subtypes involved in the cytokine storm. DC, dendritic cells. Epi, epithelial cells. Macro, macrophage cells. Mono, monocytes. Neu, neutrophils.

To examine how hyper-inflammatory cells interacted with each other in driving the inflammatory cytokine storm, we analyzed the ligand-receptor pairing patterns among hyper-inflammatory cell subtypes in severe and moderate samples within PBMC and BALF respectively (Figure S7). The interactions between PBMCs and BALF cells appeared to show significant alterations (Figure 7D). Our data revealed elevated ligand-receptor interactions of hyper-inflammatory cells in patients at severe compared to moderate stage. Interestingly, cells in the peripheral blood of severe patients showed much lower interactions with each other compared to those in BALF (Figure. S7A), except for the megakaryocytes, which secreted IL1B and stimulated Mono_c1-CD14-CCL3 cells. Mono_c1-CD14-CCL3 cells in BALF expressed *CCR5,* which could receive multiple cytokine stimulations secreted by other cell types in both the lung tissue and the peripheral blood. By contrast, the interactions of Macro_c2-CCL3L1 cells mainly relied on *CCR2* and *IL1R2.* Collectively, these findings illustrated the molecular basis for the potential cell-cell interactions at the pulmonary interface in an inflamed state, leading to a better understanding of the mechanisms of SARS-CoV-2 infection.

## DISCUSSION

Our SC4 alliance members generated scRNA-seq data for 284 clinical samples from 205 COVID-19 patients and healthy controls in China, and constructed an information-rich data resource for dissecting the immune responses of COVID-19 patients at the single-cell resolution. We observed a significant reduction of total T cells in the peripheral blood of COVID-19 patients but no notable changes of NK cells, consistent with previous observations (Liao et al., 2020). However, we did not observe a decrease of total B cells, but instead noted elevation in some patients, particularly those with severe symptoms. This contradicts previous studies based on flow cytometry (Giamarellos-Bourboulis et al., 2020), which may reflect sampling fluctuation in small cohorts instead of technology bias, although this has yet to be confirmed. Our findings indicate that T cell changes may be a major cause of the lymphopenia in COVID-19 patients. Despite the overall reduction of total T cells in the peripheral blood, proliferative CD4+ and CD8+ T cells were actually elevated in peripheral blood and were enriched in lung samples, indicating activated cellular immune responses to SARS-CoV-2 infection. Similarly, despite the conflicting reports on total B cell levels, plasma B cells were consistently elevated in patient lung samples and blood, supporting an activated humoral response (Gudbjartsson et al., 2020; Ni et al., 2020b). The complex patterns of T and B cell subtype changes indicate that additional investigations are needed to understand the detailed mechanism by which the cellular and humoral immune responses are activated and compromised in COVID-19 patients.

Previous clinical and epidemiological studies have revealed obvious sex and age biases in infection rates and disease severity of COVID-19 patients. Our data, covering a wide age range and a sex-balanced COVID-19 cohort, proved to be powerful at dissecting the associations of age and sex in the immune responses to SARS-CoV-2 infection. Our data revealed an apparent involvement of age and sex in the diverse human immune responses via multiple mechanisms, at least partially reflected at the immune cell sub-cluster level. In general, plasma B and proliferative T cells were associated with disease severity, while compositional differences of the precursor cells of these adaptive immune cell types were more prone to be influenced by sex and age seemed to impact more on naive and central memory cells. Of note, age and sex also seemed to impact the diversity of TCR/BCR repertoires for a wide range of T and B cells, which may have clinical implications.

The single-cell resolution of our data also enabled us to examine the *in vivo* potential host cells of SARS-CoV-2 and the transcriptomic changes caused by SARS-CoV-2 infection. We observed the presence of SARS-CoV-2 RNAs in multiple epithelial cell types in the human respiratory tract, including ciliated, secretory, and squamous cells. Although prominent type I interferon responses could be identified in these cells, distinct transcriptomic changes appeared to be caused by SARS-CoV-2 infection. Such distinctions were exhibited not only in the correlations of interferon responses and viral load, but also in the genes of specific immune relevance, including those encoding LR interactions which are pivotal to cell-cell communications. Of hundreds of immune-relevant LR pairs, ANXA1-FPR1 and S100A8/A9-TLR4 seemed to be critical in mediating the interactions of virus+ squamous epithelial cells and neutrophils and macrophages. Although *S100A8/A9* were not expressed in lymphoid cells in mild/moderate COVID-19 patients, they were highly expressed in the T, B, and NK cells of severe patients, likely contributing to the aberrant inflammation of these patients. Coincidentally, small molecule inhibitors of S100A8/A9 could reduce the aberrant inflammation and SARA-CoV-2 replication in mice (Guo et al., 2020b), supporting our findings. Both S100A8/9 and FPR1 should be evaluated further as targets for modulating the immune responses to SARS-CoV-2.

In addition to epithelial cells, RNAs of SARS-CoV-2 were also identified in various immune cell types, including neutrophils, macrophages, plasma B cells, T and NK cells, often with even higher levels than those in epithelial cells. The viral infection status of these cells could also be supported by the prominent interferon responses in these cells. It is still not clear how such immune cells would acquire viral sequences in the absence of either ACE2 or TMPRSS2, but it is evident that the pattern of SARS-CoV-2 infection is more complicated than initial understanding. Such complexity needs to be thoroughly addressed before this dreadful infectious disease can be effectively controlled.

The rich information of our data also allowed us to dissect the cellular origins of potential cytokine storms. We found that megakaryocytes and a few monocyte subsets might be key sources of a diverse set of cytokines highly elevated in COVID-19 patients with severe disease progression. We suspect that in severe patients, infected epithelial cells would secrete cytokines such as IL1RN into the peripheral, and monocytes expressing IL1R2 could be stimulated and in turn produce multiple proinflammatory cytokines such as CXCL8, IL6, IL1B, and TNF (Figure 7E). Through IL1R2, these hyperactive monocytes could also interact with dysfunctional megakaryocytes producing TGFB1, TNFSF4, PF4 and FTH1. Meanwhile, the T cells in the blood go through lymphopenia, while the residual ones are hyperactive in secreting many inflammatory cytokines such as IFNG and TNFSF8. Such proinflammatory cytokines secreted by the cells in the blood could also infiltrate into the lung tissue, and thus activating the tissue resident monocytes, macrophages and neutrophils for further cytokine production. We acknowledge that this is only one of many possible scenarios where an inflammatory storm could form, although our data revealed key actors in the final cytokine screenplay.

In conclusion, we generated a large scRNA-seq dataset including ~1.5 million single cells covering diverse disease severity and stages. Analyses based on this dataset revealed multiple immune characteristics of SARS-CoV-2 infection with single-cell resolution. Such data provide a critical resource and important insights in dissecting the pathogenesis of COVID-19, and potentially help the development of effective therapeutics and vaccines against SARS-CoV-2.

## ACKNOWLEDGMENTS

We thank the Computing Platform of the Center for Life Science for their contributions. We also thank 10x Genomics China team for their strong support in coordination and discussion of this study. We thank Analytical BioSciences for their help in building the visualization website of this dataset. We thank Zhuo Zhou of BIOPIC, Peking University for his helpful discussion on the interferon responses of SARS-CoV-2 infection. We thank the USTC supercomputing center and the School of Life Science Bioinformatics Center for providing supercomputing resources for this project. We thank the CAS interdisciplinary innovation team for helpful discussion. We also thank W.Chen, C.Xiao, Y.Zheng, W.Su, Y. Zhang, C. Zhang, X.Wang, H.Ma, T.Jin, X.Wang, H.Wei, B.Fu, L.Liu, J.Weng, X.Ma for their support to this project. This work was supported by the Beijing Advanced Innovation Centre for Genomics at Peking University, the Ministry of Science and Technology of the People’s Republic of China (2020YFC0848700), the National Key R&D Program of China (2017YFA0102900 to K.Q.), the National Natural Science Foundation of China grants (91940306, 81788101, 31970858, 31771428, 91640113 to K.Q., 31700796 to C.G., 81871479 to J.L., 81772165, 81974303, 31991171,31530036 and 91742203), the China Primary Health Care Foundation-Youan Medical Development Fund (BJYAYY-2020PY-01), the Beijing Municipal of Science and Technology Major Project (Z2011000054200018), the Fundamental Research Funds for the Central Universities (YD2070002019 to K.Q.).

## AUTHOR CONTRIBUTIONS

Conceptualization, Z.Z., X.R., R.J., J.C., X.W, K.Q.,Z.Z., H.W., F.W., P.Z., X.L.,T.C., X.L., L.W., J.B., Z.H., Q.J. and P.Z.; Resources, Z.Z., X.R., R.J., J.C., X.W, K.Q.,Z.Z., H.W., F.W., P.Z., X.L.,T.C., X.L., L.W., J.B., Z.H., Q.J. and P.Z.; Methodology, Z.Z., X.R., R.J., J.C., X.W, K.Q.,Z.Z., H.W., F.W., P.Z., X.L.,T.C., X.L., L.W., J.B., Z.H., Q.J., P.Z., W.W., X.F., W.H., B.S., P.C., J.L., Y.L., F.T., F.Z., Y.Y., J.H., W.M., X.X. and P.W.; Investigation, Z.Z., X.R., R.J., J.C., X.W, K.Q.,Z.Z., H.W., F.W., P.Z., X.L.,T.C., X.L., L.W., J.B., Z.H., Q.J., P.Z., W.W., X.F., W.H., B.S., P.C., J.L., Y.L., F.T., F.Z., Y.Y., J.H., W.M., X.X. and P.W.; Validation, W.W. and W.Y.; Writing Original Draft, Z.Z., X.R., R.J., J.C., X.W, K.Q.,Z.Z., H.W., F.W., P.Z., X.L.,T.C., X.L., L.W., J.B., Z.H., Q.J., P.Z., W.W., X.F., W.H., B.S., P.C., J.L., Y.L., F.T., F.Z., Y.Y., J.H., W.M., X.X. and P.W.

## DECLARATION OF INTERESTS

Z.Z. is a founder of Analytical Bioscience and an advisor for InnoCare. All financial interests are unrelated to this study. Other authors declare no competing interests.

## STAR+METHODS

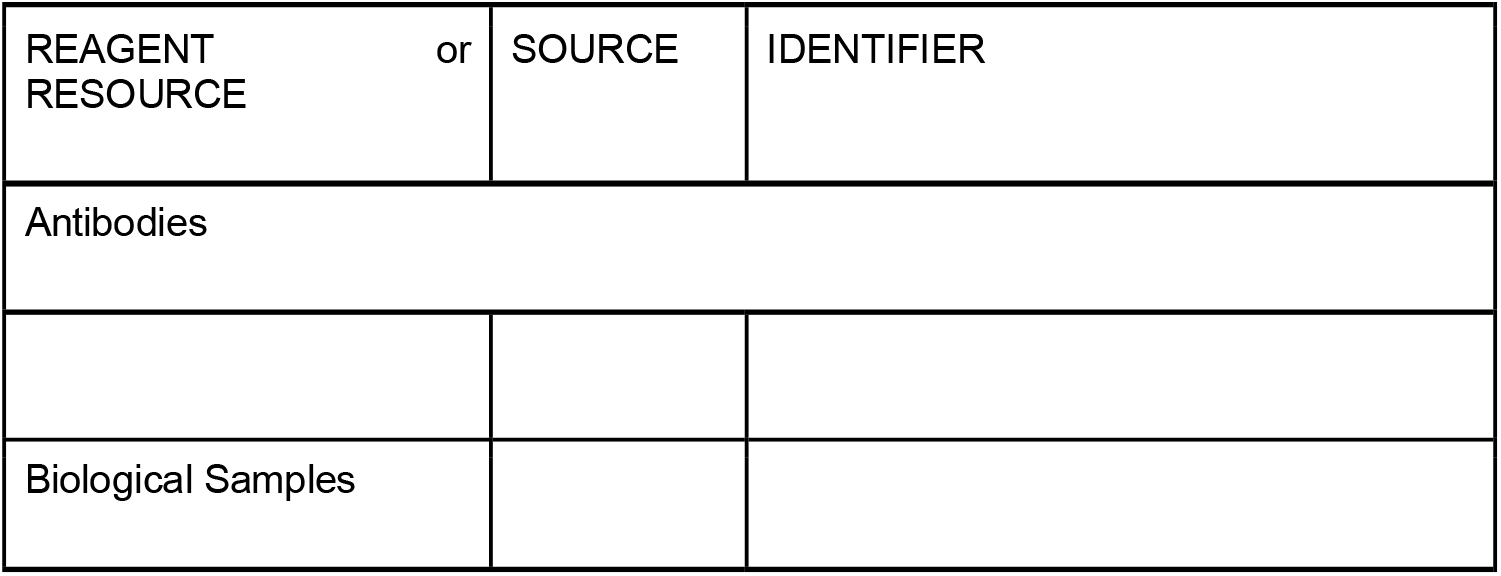
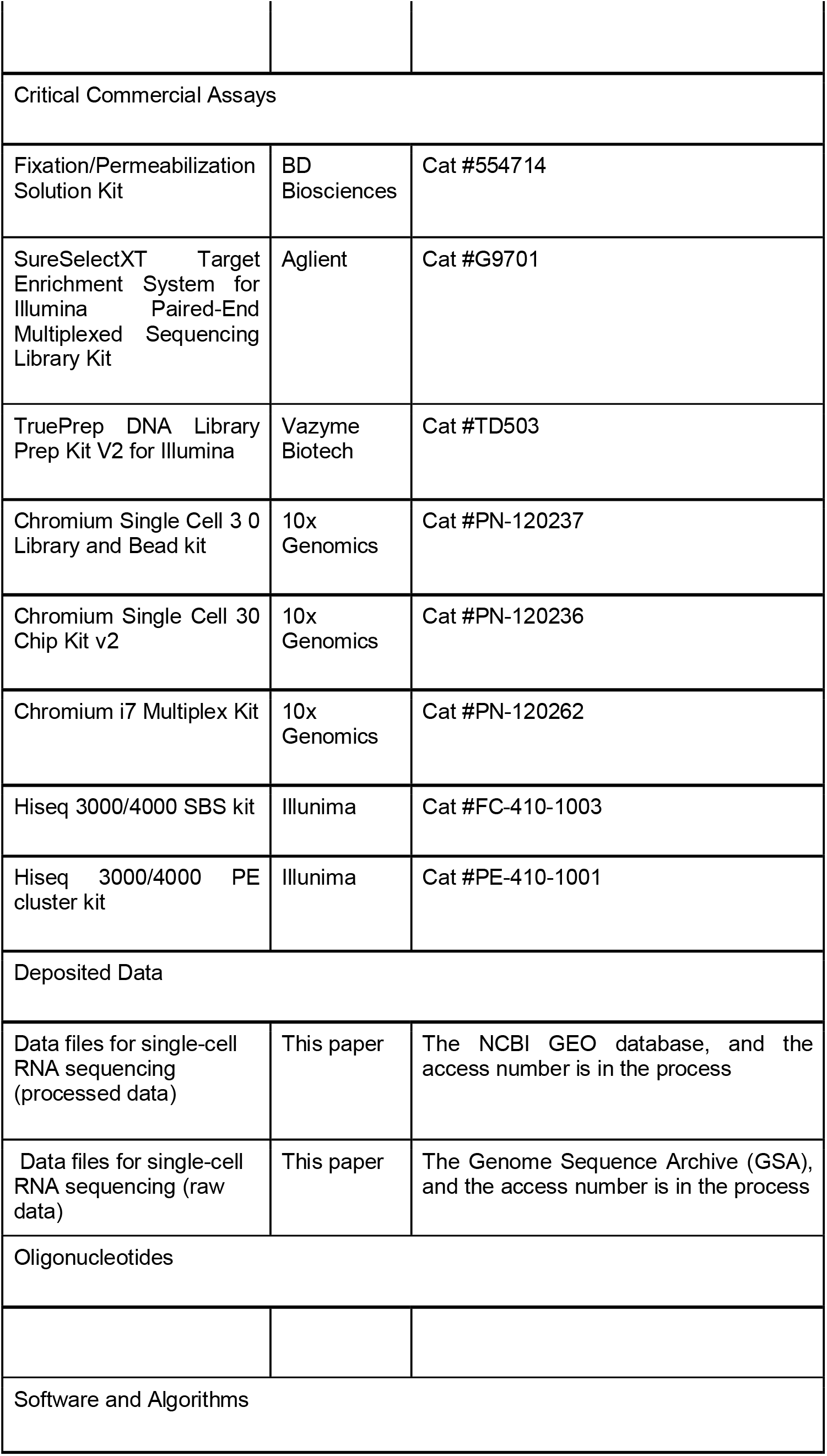
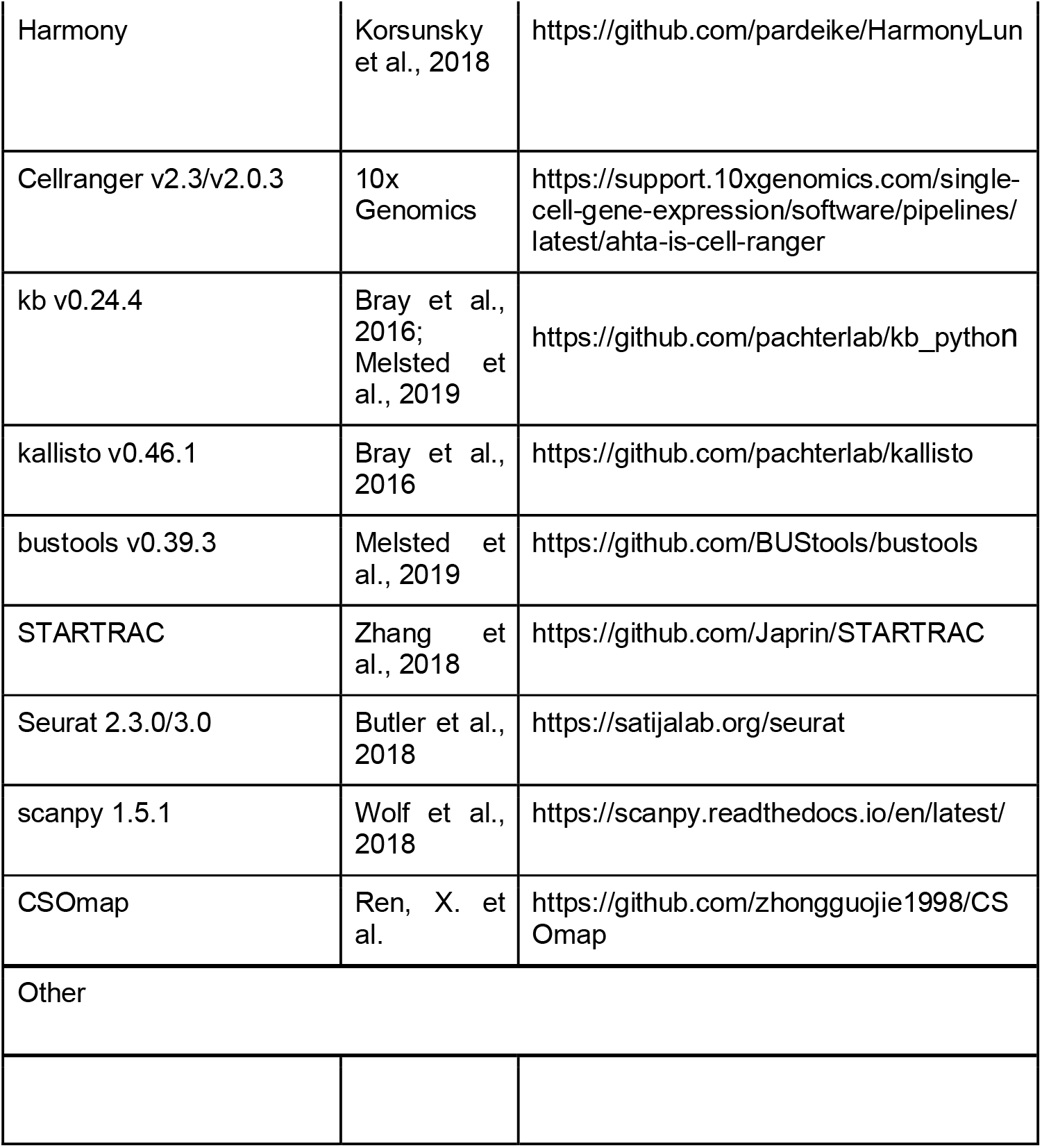
KEY RESOURCES TABLE

## LEAD CONTACT AND MATERIALS AVAILABILITY

Further information and requests for resources and reagents should be directed to and will be fulfilled by the Lead Contact, Zemin Zhang.

## EXPERIMENTAL MODEL AND SUBJECT DETAILS

### Ethics statement

This study was approved by the Ethics Committees of respective institutions, with written informed consents obtained from all participants before sample collection according to regular principles.

### Human subjects

A total of 183 patients with COVID-19 and 25 healthy individuals in this study were enrolled from 36 centers/ laboratories, with samples (n=284) collected. Samples of COVID-19 were further categorized into groups of moderate convalescence (n=83), moderate progression (n=33), severe convalescence (n=51) and severe progression (n=83) according to disease severity (moderate or severe) and stages (progression and convalescence) based on the Guidelines for Diagnosis and Treatment of Corona Virus Disease 2019 issued by the National Health Commission of China (7th edition). The sex ratio between female and male donor is 101:159. The age of the donors ranges from 6 to 92. Of all the 284 samples, 249 samples were collected from PBMC, among which 77 samples have sorted B/T cells or both. 13 samples were collected from lung tissues, including 12 BALF samples and 1 PFMC sample. We also collected 22 sputum samples from patients as well. Among all the samples, we have 7 paired lung BALF and PBMC samples. Single-cell transcriptome data for each sample was profiled using 10x Genomics scRNA-seq platform. Single-cell sequencing of TCRs (13 samples) and BCRs (53 samples) or both (11 samples) was also performed for part of the samples. Detailed clinical information and demographic characteristics of patient cohorts were shown in Table S1.

## METHOD DETAILS

### Sample collection

Blood samples that were not immediately processed for cell encapsulation were mixed with Whole Blood Cell Stabilizer (Cytodelics) and stored at −80 °C freezer. The peripheral blood mononuclear cell (PBMCs) were isolated using standard density gradient centrifugation and then used for 10x single-cell RNA-seq. Bronchoalveolar lavage fluid (BALF) samples were collected from COVID-19 patients and processed with 2h according to WHO guidance. BALF was passed through 100-μm nylon cell strainer to obtain single cell suspensions with cooled RPMI 1640 complete medium. Cells in the BALF were freshly used for 10x singlecell RNA-seq. Sputum samples were collected from COVID-19 patients using an oropharyngeal swab or hypertonic saline induction. To reduce squamous cell contamination, subjects were asked to rinse their mouth with water and clear their throat. Samples were incubated in Dulbecco’s Phosphate-Buffered Saline (DPBS) with agitation for 15 minutes and filtered through 40-micron strainers. Cells in the sputum were freshly used for 10x single-cell RNA-seq.

### Single cell RNA library preparation and sequencing

Cell suspensions were barcoded through the 10x Chromium Single Cell platform using Chromium Single Cell 5’ Library, Chromium Single Cell 3’ Library, Gel Bead and Multiplex Kit, and Chip Kit (10x Genomics). The loaded cell numbers range from 300-500,000 aiming for 300-14,000 single cells per reaction. Single-cell RNA libraries were prepared using the Chromium Single Cell 3’ v2 Reagent (10x Genomics; PN-120237, PN-120236 and PN-120262), Chromium Single Cell 3’ v3 Reagent (10x Genomics; PN-1000075, PN-1000073 and PN-120262) the Chromium Single Cell 5’ v2 Reagent (10x Genomics, 120237), and Chromium Single Cell V(D)J Reagent kits (10x Genomics, PN-1000006, PN-1000014, PN-1000020, PN-1000005) was used to prepare single-cell RNA libraries according to the manufacturer’s instructions. Each sequencing library was generated with a unique sample index. The libraries were sequenced using either DIPSEQ, BGISEQ or Illumina platforms.

### Single-cell RNA-seq data processing

Single-cell sequencing data were aligned and quantified using kallisto/bustools (KB, v0.24.4) (Bray et al., 2016) against the GRCh38 human reference genome downloaded from 10x Genomics official website. Preliminary counts were then used for downstream analysis. Quality control was applied to cells based on three metrics step by step: the total UMI counts, number of detected genes and proportion of mitochondrial gene counts per cell. Specifically, cells with less than 1000 UMI counts and 500 detected genes were filtered, as well as cells with more than 10% mitochondrial gene counts. To remove potential doublets, for PBMC samples, cells with UMI counts above 25,000 and detected genes above 5,000 are filtered out. For other tissues, cells with UMI counts above 70,000 and detected genes above 7,500 are filtered out. Additionally, we applied Scrublet (Wolock et al., 2019) to identify potential doublets. The doublet score for each single cell and the threshold based on the bimodal distribution was calculated using default parameters. The expected doublet rate was set to be 0.08, and cells predicted to be doublets or with doubletScore larger than 0.25 were filtered. After quality control, a total of 1,598,708 cells were remained. We normalized the UMI counts with the deconvolution strategy implemented in the R package scran (Lun et al., 2016). Specifically, cell-specific size factors were computed by computeSumFactors function and further used to scale the counts for each cell. Then the logarithmic normalized counts were used for the downstream analysis.

### Batch effect correction and cell subsets annotations

To integrate cells into a shared space from different datasets for unsupervised clustering, we used the harmony algorithm (Korsunsky et al., 2019) to do batch effect correction. To detect the most variable genes used for harmony algorithm, we performed variable gene selection separately for each sample. A consensus list of 1,500 variable genes was then formed by selecting the genes with the greatest recovery rates across samples, with ties broken by random sampling. All ribosomal, mitochondrial and immunoglobulin genes were then removed from the list. Next, we calculate a PCA matrix with 20 components using such informative genes and then feed this PCA matrix into HarmonyMatrix() function implemented in R package Harmony. We set sample and dataset as two technical covariates for correction with theta set as 2.5 and 1.5, respectively. The resulting batch-corrected matrix was used to build nearest neighbor graph using scanpy (Wolf et al., 2018). Such nearest neighbor graph was then used to find clusters by Louvain algorithm (Traag et al., 2019). The cluster-specific marker genes were identified using the rank_genes_groups function.

The first round of clustering (resolution = 0.3) identified six major cell types including T cells, NK cells, B cells, plasma B cells, myeloid cells and epithelial cells. To identify clusters within each major cell type, we performed a second round of clustering on T/NK, B/plasma B, myeloid and epithelial cells separately. The procedure of the second round of clustering is the same as first round, starting from low-rank harmony output (30 components) on the highly variable genes chosen as described above, with resolution ranging from 0.3 to 1.5. Each sub cluster was restrained to have at least 30 significantly highly expressed genes (FDR < 0.01, logFC > 0.25, t test) compared with other cells. Annotation of the resulting clusters to cell types was based on the known markers. Meanwhile, single cells expressing two sets of well-studied canonical markers of major cell types were labeled as doublets and excluded from the following analysis. Also, cells highly expressed *HBA, HBB* and *HBD*, which are the markers for erythrocytes, were also excluded. 136,006 cells were removed and a total of 1,462,702 cells were retained for downstream analysis.

### Detection and processing of cells with viral RNA

To identify single cells with viral infection, we aligned raw scRNA-seq reads using kallisto/bustools(KB) against a customized reference genome, in which the SARS-CoV-2 genome (Refseq-ID:NC_045512) was added as an additional chromosome to the human reference genome. Single cells with viral reads (UMI > 0) were retained. Cells with less than 200 genes expressed or more than 20% mitochondrial counts were excluded, as well as those labeled as doublet following aforementioned protocol.

The remaining cells were then used for dimension reduction and unsupervised clustering using Python package scanpy (Wolf et al., 2018) In brief, the top 500 genes with the highest variance were selected and the dimensionality of the data was reduced by principal component analysis (PCA) (30 components) first and then with t-SNE, followed by Louvain clustering (Traag et al., 2019) performed on the 30 principal components (resolution = 1). For t-SNE visualization, we directly fit the PCA matrix into the scanpy.api.tl.tsne function with perplexity of 30. To identify cell-type-specific gene markers, we selected genes that were differentially expressed across different cell types (FDR < 0.01, log fold change > 0.5) using the rank_genes_groups function. Clusters were annotated based on the expression of known marker genes.

### TCR and BCR analysis

TCR/BCR sequences were assembled and quantified following Cell Ranger (v.3.0.2) vdj protocol against GRCh38 reference genome. Assembled contigs labeled as low-confidence, non-productive or with UMIs < 2 were discarded.

To identify TCR clonotype for each T cell, only cells with at least one TCR α-chain (TRA) and one TCR β-chain (TRB) were remained. For a given T cell, if there are two or more α or β chains assembled, the highest expression level (UMI or reads) α or β chains was regarded as the dominated α or β chain in the cell. Each unique dominated α-β pair (CDR3 nucleotide sequences and rearranged VDJ genes included) was defined as a clonotype. T cells with exactly the same clonotype constituted a T cell clone.

BCR clonotypes were identified similar to TCR. Only cells with at least one heavy chain (IGH) and one light chain (IGL or IGK) were kept. For a given B cell, if there are two or more IGH or IGL/IGK assembled, the highest expression level (UMI or reads) IGH or IGL/IGK was defined as the dominated IGH or IGL/IGK in the cell. Each unique dominated IGH-IGL/IGK pair (CDR3 nucleotide sequences and rearranged VDJ genes) was defined as a clonotype. B cells with exactly the same clonotype constituted a B cell clone.

220,968 T cells with TCR information and 282,464 B cells with BCR information were used to perform the STARTRAC analysis as we previously described (Zhang et al., 2018).STARTRAC-expa was used to quantified the potential clonal expansion level. TCR/BCR diversity was calculated as Shannon’s entropy shown below:

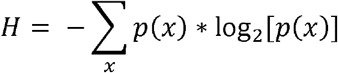

The *p*(x) represents the frequency of a given TCR/BCR clone among all T/B cells with TCR/BCR identified.

### Comparing immune cell proportion

For samples from PBMC and BALF tissue, we calculated immune cell proportions for each major cell type and underlying cell subsets. In order to avoid bias caused by samples dominated by few cell types, we filtered samples containing FACS-sorted B/T cells and retained those samples with CD45+ cells > 1000. For each sample, cell type proportion was calculated by number of cells in certain cell type divided by total number of CD45+ cells. To identify changes in cell proportions between samples in different disease severity states, disease progression stages and sex, we performed Wilcoxon rank-sum test on the proportions of each major cell type and underlying cell subset across different phenotype groups. We performed correlation analysis to assess the association between cell type proportion and patient age. Only those cell types with statistically significant differences (FDR < 0.05) in proportions were shown.

### ANOVA analysis

To further assess how different patients’ phenotypes and their potential interactions influence cell type proportions, we performed multivariate ANOVA on cell type proportions and on diversity of BCR/TCR based on different patient phenotypes, including disease severity, disease progression stage, sex and age. All interactions between these variables were included in the models. To convert age into a categorical variable, we binned patient age into four groups: young (<18 years old), middle-age (18-50 years old), old-age (50-70 years old) and the elderly (70+ years old). Interactions between variables were regarded as significantly associated with cell type proportions when FDR < 0.05.

### Differential expression and GO term enrichment analysis

To investigate the impact of virus infection on epithelial cells, we identify differential expressed genes by performing two-sided unpaired Wilcoxon tests on all the expressed genes (expressed in at least 10% cells in either group of cells). P values were adjusted following Benjamini & Hochberg protocol. Top 100 highly expressed genes of each group were shown in the volcano plots. Based on these genes, enriched GO terms were then acquired for each group of cells using R package clusterProfiler (Yu et al., 2012).

### Cell-cell interaction analysis by CSOmap

To illustrate the cell-cell interaction potential of cells with viral detection, we first created a set of datasets by joining 7 BALF samples with the virus+ dataset separately. Then, we used CSOmap (Ren et al., 2020) to construct a 3D pseudo space and calculate the significant interaction for each dataset. To investigate the interaction potentials of the cell types, we used two indexes, distances within cell type and normalized connection. Distance within each cell type is calculated based on the aforementioned 3D coordinates. The shorter the distance, the closer the cells are located in the 3D space, which indicates that they are more likely to interact with each other. To further investigate the interaction between different cell types, we made use of the CSOmap output connection matrix. For a cluster pair, normalized connection was calculated by dividing its corresponding connection value by the product of their respective cell numbers. Normalized connections were then multiplied by 10,000. Meanwhile, to highlight the key ligand-receptor pairs function in the interaction, we also examine the contribution output by CSOmap.

In addition, normalized connections were also calculated on another set of cohorts where we combined virus+ dataset with samples with paired PBMC and BALF tissues, in order to investigate the interaction potential between cells from two tissues, PBMC and BALF.

### Inflammatory and cytokine score related subtypes analysis

Briefly, we firstly filtered out samples with less than 1000 cells. For PBMC, only subtypes with more than 1000 cells were included in the subsequent analysis. For BALF data analysis, we removed major cell types with less than 500 cells. To define inflammatory and cytokine score, we downloaded a gene set termed ‘HALLMARK_INFLAMMATORY_RESPONSE’ from MSigDB (PMID: 26771021) and collected cytokine genes based on these references (see Table S1). Cytokine and inflammatory score were evaluated with the AddModuleScore function built in the Seurat (PMID: 31178118). To select the most promising hyper-inflammatory cell types, we performed Mann-Whitney rank test (single-tail) for each subtype’s score versus all the other subtypes’ score. Seven subtypes (Mono_c1-CD14-CCL3, Mono_c2-CD14-HLA-DPB1, Mono_c3-CD14-VCAN, T_CD4_c08-GZMK-FOS^high^, T_CD8_c06-TNF, T_CD8_c09-SLC4A10 and Mega) in PBMC were defined as hyper-inflammatory cell types with significantly statistical parameters *(P* < 0.0001) in both cytokine and inflammatory score. In addition, we defined 8 subtypes (T_CD8_c08-IL2RB, T_CD4_c11-GNLY, NK_c01-FCGR3A, T_CD8_c05-ZNF683, T_CD8_c04-COTL1, T_CD8_c07-TYROBP, T_CD8_c03-GZMK and T_gdT_c14-TRDV2) with significantly statistical parameters (*P* < 0.0001) only in cytokine score. For subtypes in BALF, we defined 5 subtypes (Macro_c2-CCL3L1, Mono_c1-CD14-CCL3, Mono_c2-CD14-HLA-DPB1, Mono_c3-CD14-VCAN, Neu) as hyper-inflammatory types with the same standard threshold as PBMC (*P* < 0.0001).

### Cell ratio and cytokine marker analysis of hyper-inflammatory and cytokine subtypes (integrated with Statistics section)

To explore whether there are state-specific of COVID-19 patients enriched subtypes, we performed hierarchical clustering with setting standard scale (0-1) for 7 hyper-inflammatory and 8 cytokine subtypes respectively. Then, we used the Wilcoxon rank-sum test to calculate the significance of cell proportion of each subtype in states (moderate convalescent, moderate progression, severe convalescent, severe progression) compared with healthy control. We also applied the ordinary least square method to calculate the correlation between age and cell proportion in different states of COVID-19 patients. For the significance of cytokine expression level with state and age, we performed Wilcoxon ranksum test and ordinary least square to assess the *P* values.

### Cell-cell communication analysis between PBMC and BALF by iTALK

To identify and visualize the possible cell-cell interactions in terms of cytokine storm between the highly inflammation-correlated cell types evaluated by the inflammation score within each tissue and the crosstalk between lung and circulating blood, we employed an R package iTALK introduced by Wang et al. (Wang et al., 2019, bioRxiv, https://www.biorxiv.org/content/10.1101/507871v1). Cytokine/chemokine category (n = 320) in the ligand-receptor database was selectively used for our purpose. Wilcoxon rank sum test was used to identify the differentially expressed genes (DEGs) between severe onset and moderate onset patient groups for each cell type. DEGs were then matched and paired against the ligand-receptor database to construct a putative cell-cell communication network. An interaction score defined as the product of the log fold change of ligand and receptor was used to rank these interactions. In addition, the expression level of both ligand and receptor were also considered. We defined severe gained interaction if a ligand gene was upregulated in severe onset group and its paired gene upregulated or remains no change. We defined severe lost interaction if a ligand(receptor) gene was downregulated in severe onset group regardless of the expression level of its paired gene.

### Cytokine analysis of serum by using multiplex bead-based immunoassay

Human cytokines in the serum were measured by Bio-plex Pro TM Human Cytokine Screening 48 plex Bio-PlexTM 200 System (# 12007283, Bio-Rad, US) and Bio-PlexTM 200 System (Bio-Rad). Bio-Plex ProTM assays are essentially immunoassays formatted on magnetic beads and are built upon three core elements of xMAP technology, fluorescently dyed microspheres (also called beads), a dedicated flow cytometer with two lasers to measure the different molecules bound to the surface of the beads, and a high-speed digital signal processor that efficiently manages the fluorescence data.

#### Sample preparation

Whole blood from COVID-19 patients and healthy controls were drawn into collection tubes containing anticoagulant. Centrifugation the tubes at 1,000 x g for 15 min at 4°C and transfer the serum to a clean polyprophylene tube, followed by another centrifugation at 10,000 x g for 10 min at 4°C to completely remove platelets and precipitates. Dilute samples fourfold (1:4) by adding 1 volume of sample to 3 volumes of Bio-Plex sample diluent. Fifty microliter of each sample were used to assay.

#### Preparation of oupled beads

Coupled beads were diluted to a 1x concentration according to the instruction. Required volume of Bio-Plex assay buffer was added to a 15 ml polypropylene tube. Vortex the stock coupled beads at medium speed for 30 sec. Carefully open the cap and pipet any liquid trapped in the cap back into the tube. Dilute coupled beads to 1x by pipetting the required volume into the 15 ml tube and vortex. Protect the beads from light with aluminum foil and equilibrate to room temperature prior to use.

#### Assay running

Add coupled beads, standards and samples to each well of the assay plate. Cover plate with a new sheet of sealing tape and protect from light with aluminum foil. Incubate on shaker at 850 ? 50 rpm at room temperature (RT). While the samples were incubating, calculate and prepare the volume of detection antibodies and detection antibody diluent needed. After washing the plate three times with 100 μl wash buffer, transfer 25 μl detection antibodies to each well using a multichannel pipet. Cover plate with a new sheet of sealing tape and protect from light with aluminum foil. Incubate on shaker at 850 ± 50 rpm for 30 min at room temperature.

#### Read plate and calculation

Bio-Plex ManagerTM software was used for data acquisition and analysis.

## DATA AND CODE AVAILABILITY

The raw sequencing and processed gene expression data in this paper have been deposited into GSA (Genome Sequence Archive in BIG Data Center, Beijing Institute of Genomics, Chinese Academy of Sciences) and the NCBI GEO database, respectively. Visualization of this dataset can be found at http://covid19.cancer-pku.cn.

## Supplementary Figures

**Figure S1.**
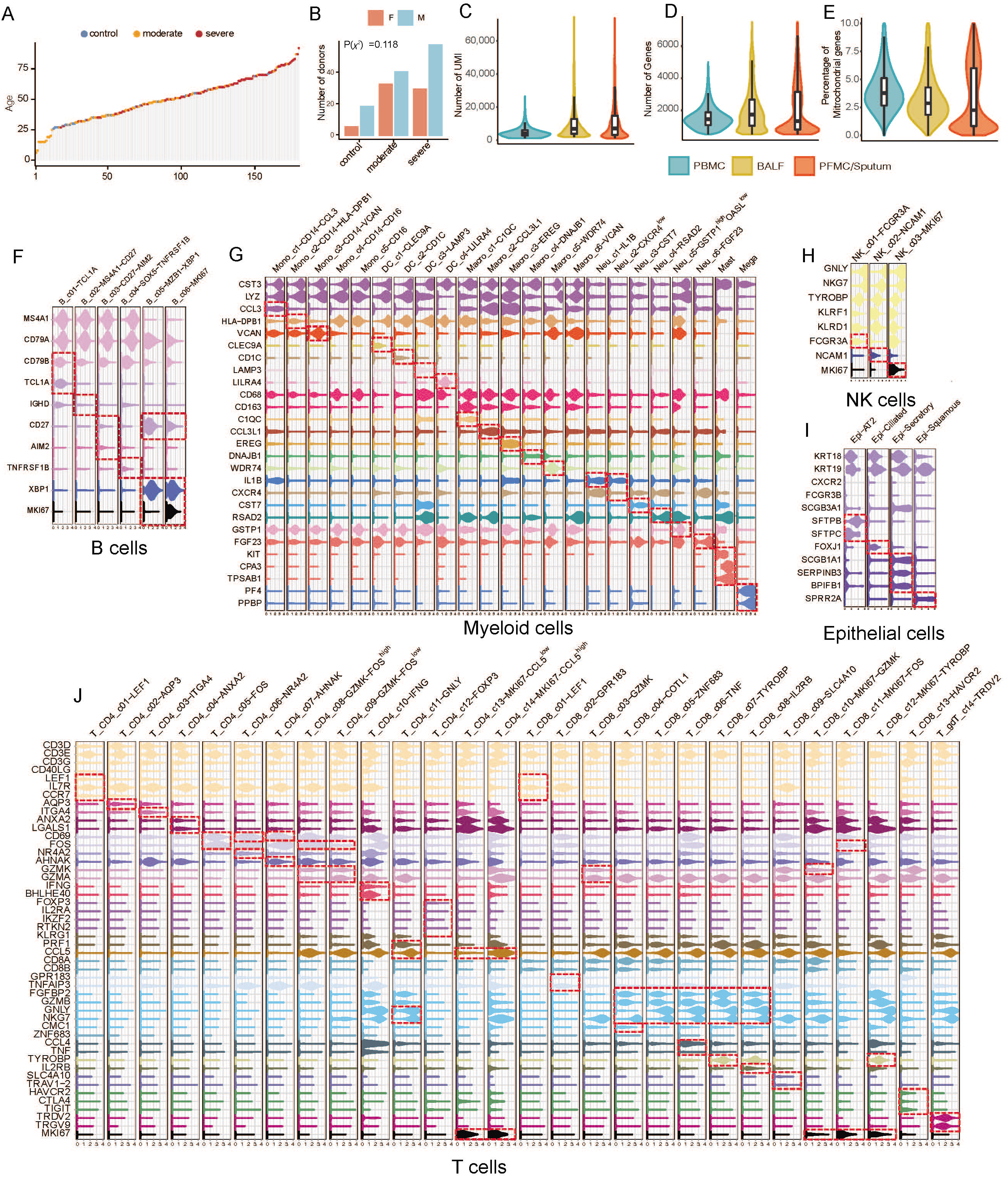
Basic information of the dataset quality and cell subsets in major cell lineages, Related to Figure 1. (A) Sorted age span of donors color-coded by disease symptoms. (B) Distribution of sex in donors with different disease symptoms. Chi-square test. (C-E) Distribution of unique molecular identifier (UMI) count per cell (C), gene count per cell (D), and percentage of mitochondrial transcripts per cell (E) detected for cells in various tissue types. PBMC, peripheral blood mononuclear cells; BALF, bronchoalveolar lavage fluid; PFMC/Sputum, pleural effusion/sputum. (F-J) Violin plots of selected marker genes (rows) for cell subsets (columns) within each cell lineage, including 6 B/plasma B cell clusters (F), 23 Myeloid cell clusters (G), 3 NK cell clusters (H), 4 Epithelial cell clusters (I) and 28 T cell clusters (J).

**Figure S2.**
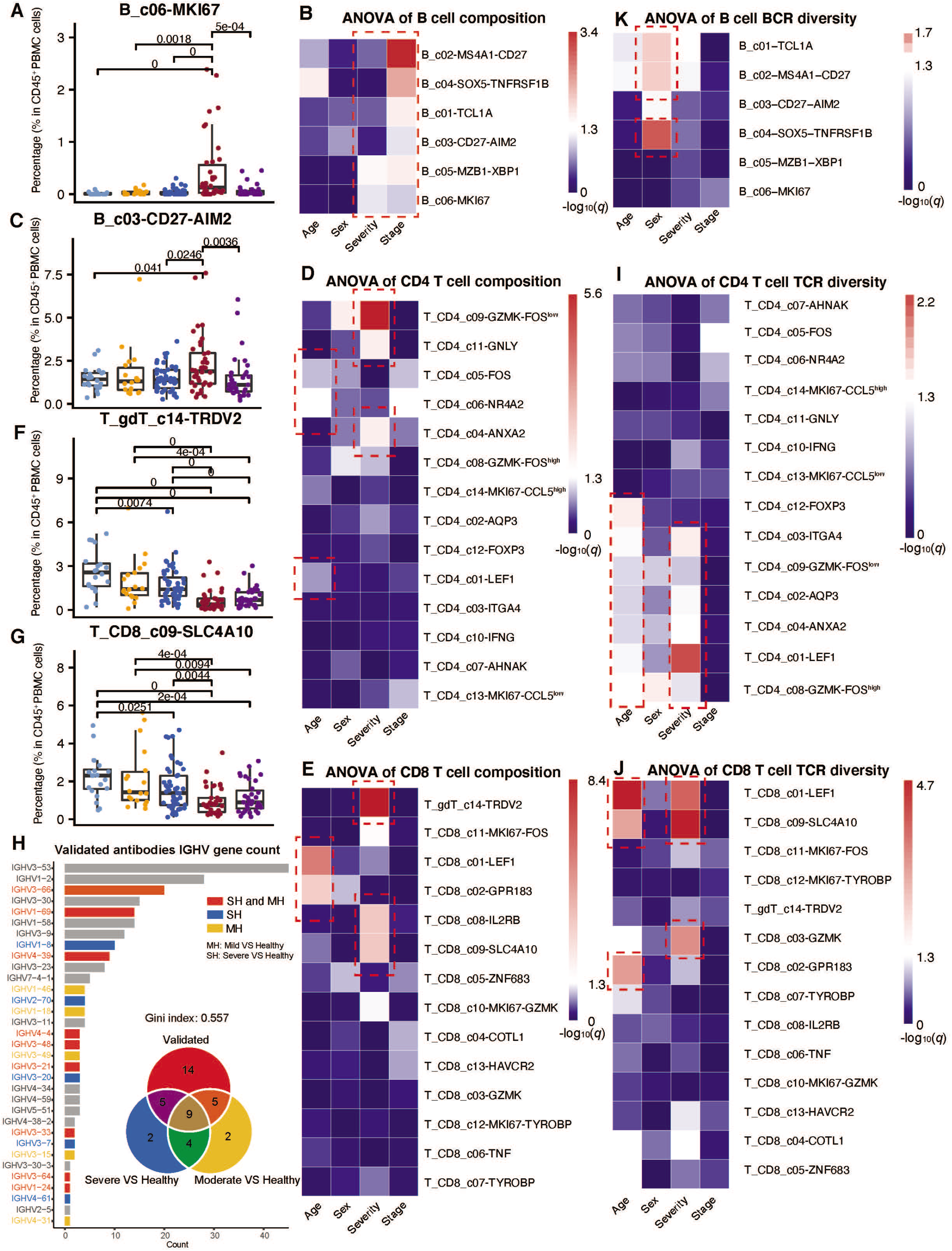
ANOVA of cell composition and clonal expansion, Related to Figures 2 and 3. (A) Differences of B_c06-MKI67 cells proportion across disease conditions. All differences with *P*-value < 0.05 are indicated; two-sided unpaired Wilcoxon test. (B) ANOVA of B cells proportion (C) Differences of B_c03-CD27-AIM2 cells proportion across disease conditions. All differences with *P*-value < 0.05 are indicated; two-sided unpaired Wilcoxon test. (D) ANOVA of CD4 T cells proportion (E) ANOVA of CD8 T cells proportion (F) Differences of T_c14_gdT-TRDV2 cells proportion across disease conditions. All differences with *P*-value < 0.05 are indicated; two-sided unpaired Wilcoxon test. (G) Differences of T_c09_CD8-SLC4A10 cells proportion across disease conditions. All differences with *P*-value << 0.05 are indicated; two-sided unpaired Wilcoxon test. (H) V gene usage of SARS-Cov-2 neutralized antibodies. (I-K) ANOVA of the diversity of TCR/BCR repertoire, estimated by Shannon’s entropy.

**Figure S3.**
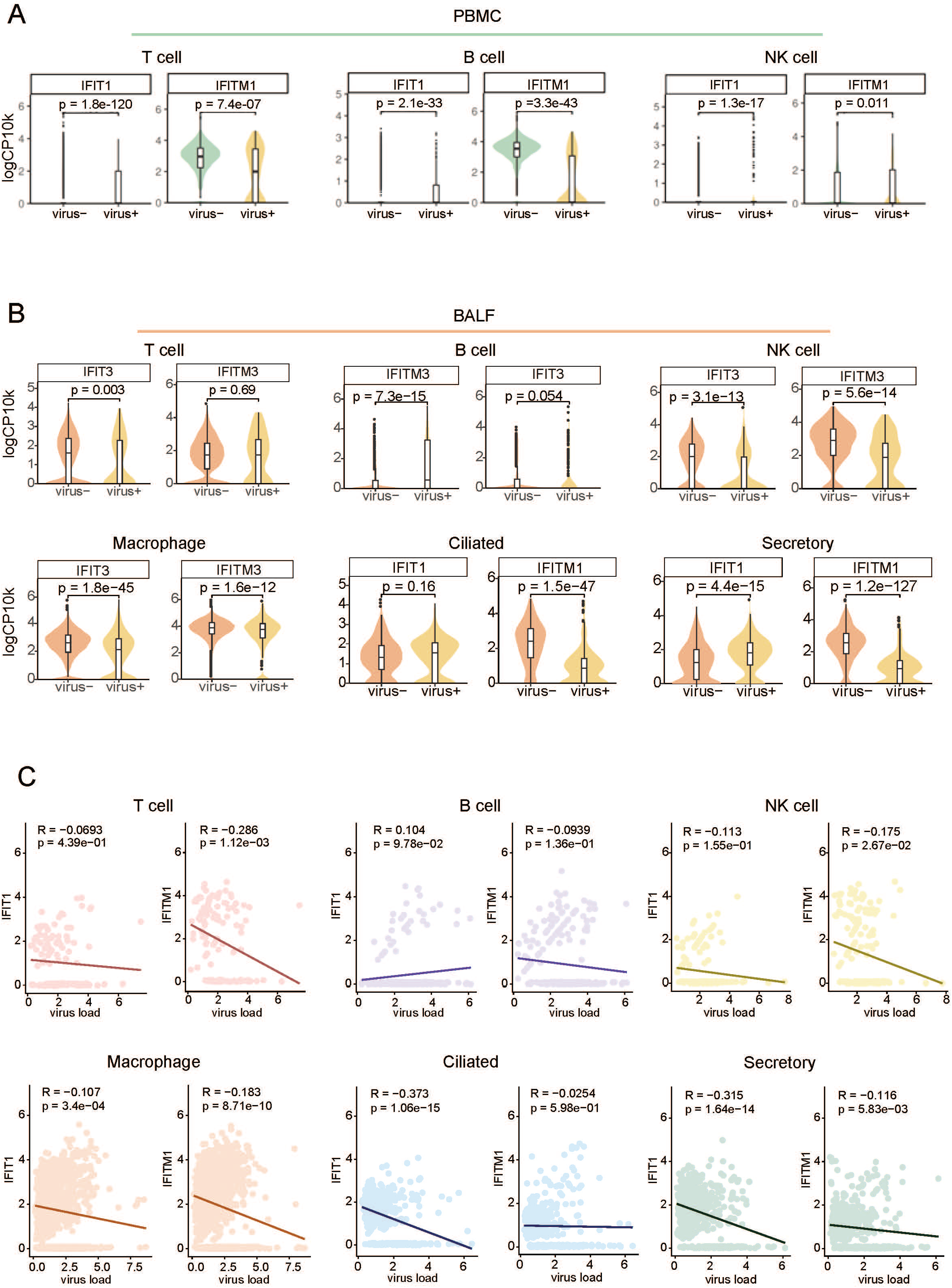
Differential Expression Analysis for ISG Genes between other Virus+ and Virus-cells in PBMC or BALF, Related to Figures 4. (A) The ISG genes in virus+ differentially expressed comparison against virus-in residential cell types including immune cells and epithelial cells. Two sided unpaired Wilcoxon test. (B) Violin plots showing the expression of ISGs in BALF. Two sided unpaired Wilcoxon test. (C) Scatter plots showing the correlation between viral load and expression level of viral load. Pearson’s correlation.

**Figure S4.**
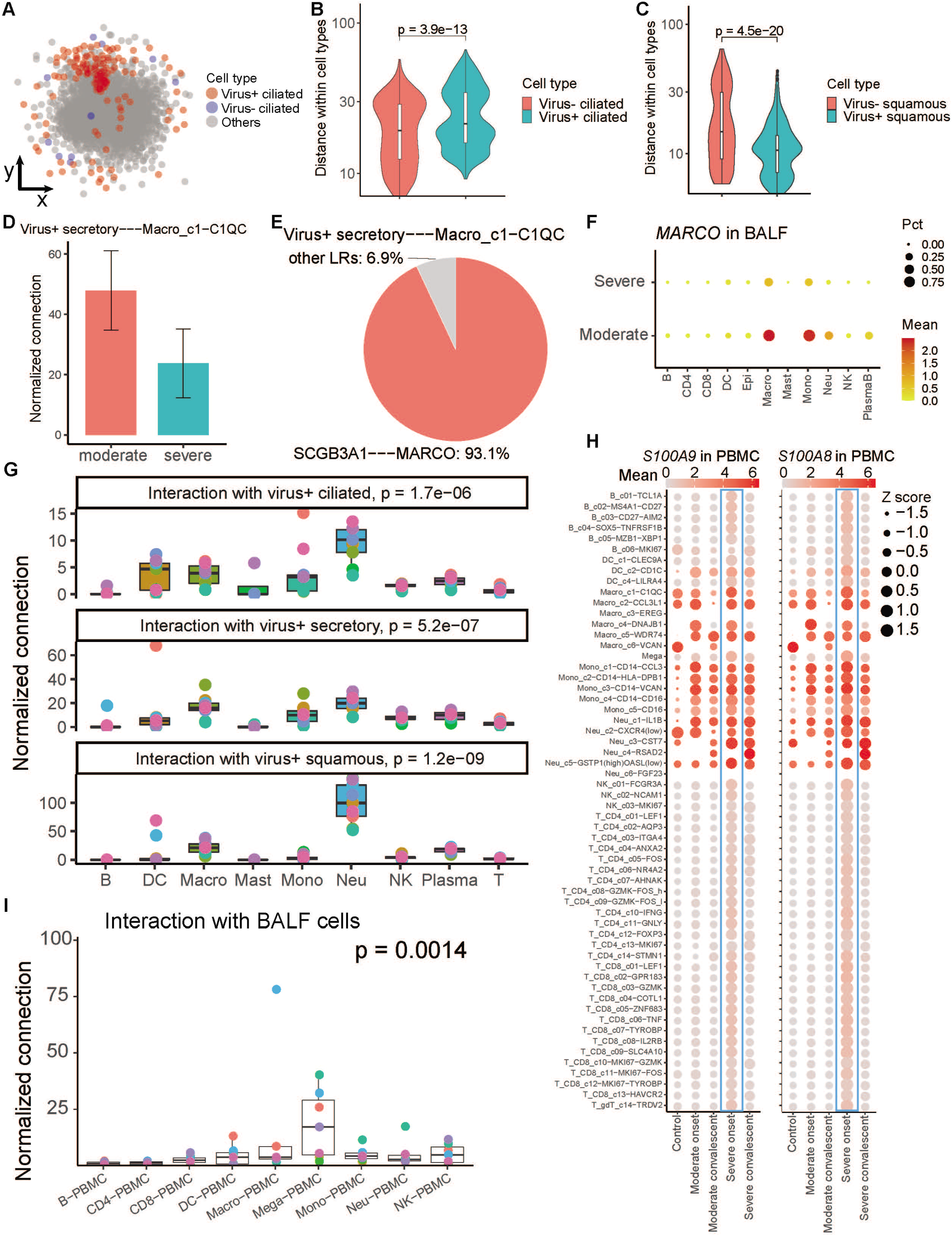
Detailed Investigation of interacting potentials of epithelial cells with viral detection, Related to Figure 5. (A) 2D pseudo space calculated by CSOmap, showing the location of ciliated cells. Each dot denotes a single cell, colored by cell type. (B) Violin plot showing the distance calculated from space shown in (A) within each ciliated cell group. Two-sided unpaired Wilcoxon test. (C) Violin plot showing the distance within each squamous cell group. Two-sided unpaired Wilcoxon test. (D) Bar plot showing the mean of normalized connections of the interaction between virus+ secretory and Macro_c1-C1QC in patients categorized by two states. Error bar, s.e.m. (E) Pie chart showing the ligand-receptor contribution proportion between virus+ secretory and Macro_c6-VCAN in one example. Ligand-receptor pairs with contribution less than 0.05 were merged as ‘Other LRs’. (F) Dotplot showing the mean expression level of *MARCO* in BALF samples. Pct, percentage of expressed cells. (G) Boxplot of normalized connection between major cell types and ciliated (top), secretory (middle) and squamous (bottom) cells with viral detection. Kruskal-Wallis Rank Sum Test. (H) Dot plots showing the expression of *S100A9*(left) and *S100A8* (right) in PBMC samples. Each dot is colored by the means of the expression and sized by the scaled means (Z scores). Blue boxes highlight expressions in severe onset patients. (I) Boxplot of normalized connection between PBMC-derived cell types and BALF. Each dot represents a sample. Kruskal-Wallis Rank Sum Test.

**Figure S5.**
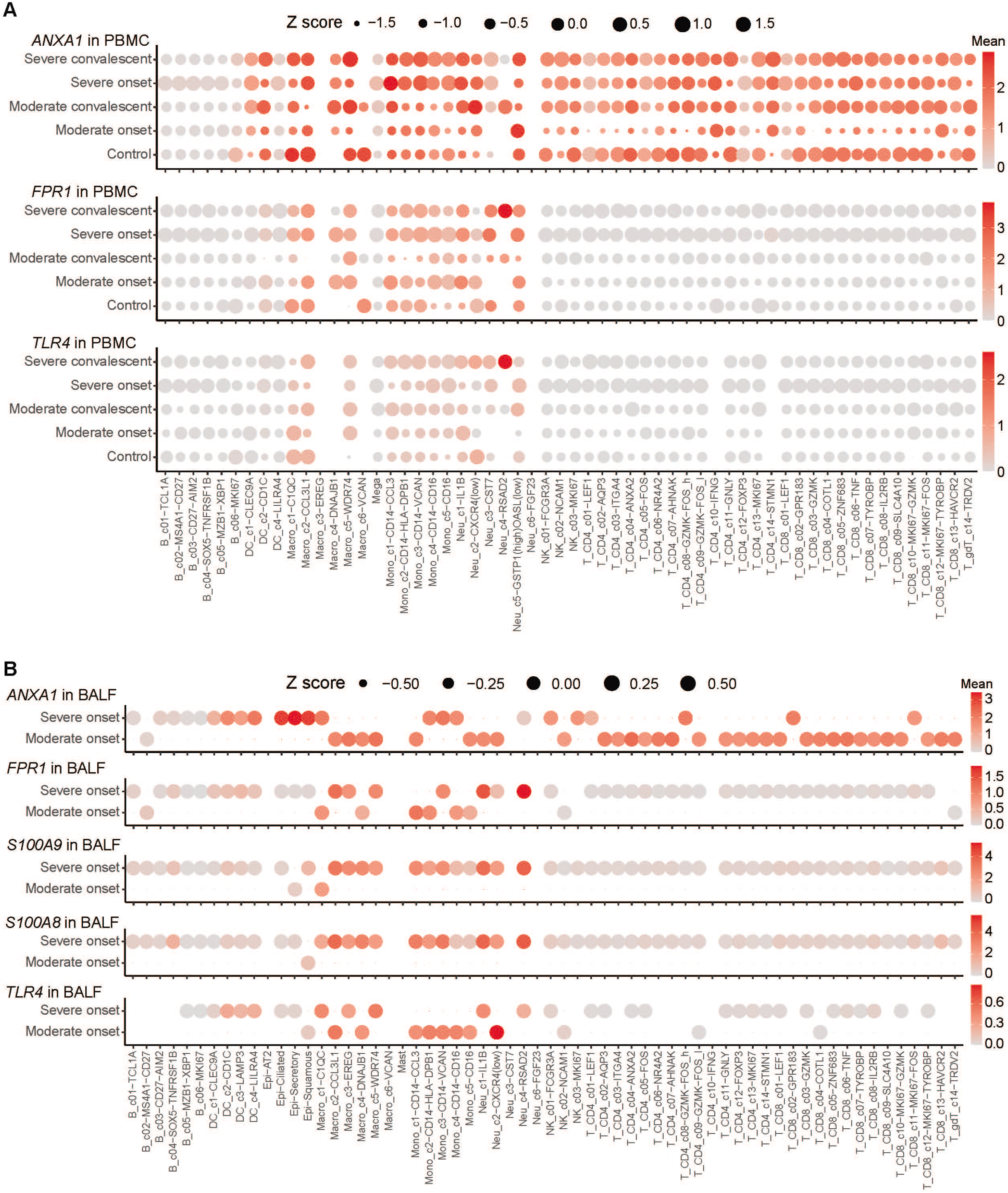
The Expression of Genes in PBMC and BALF samples, Related to Figure 5. (A) Dot plots showing the expression of *ANXA1* (top), *FPR1* (middle) and *TLR4* (bottom) in PBMC samples. (B) Dot plots showing the expression of *ANXA1* (first panel), *FPR1* (second panel) *S100A9* (third panel), *S100A8* (fourth panel) and *TLR4* (bottom panel) in BALF samples. Each dot is colored by the means of the expression and sized by the scaled means (Z scores).

**Figure S6.**
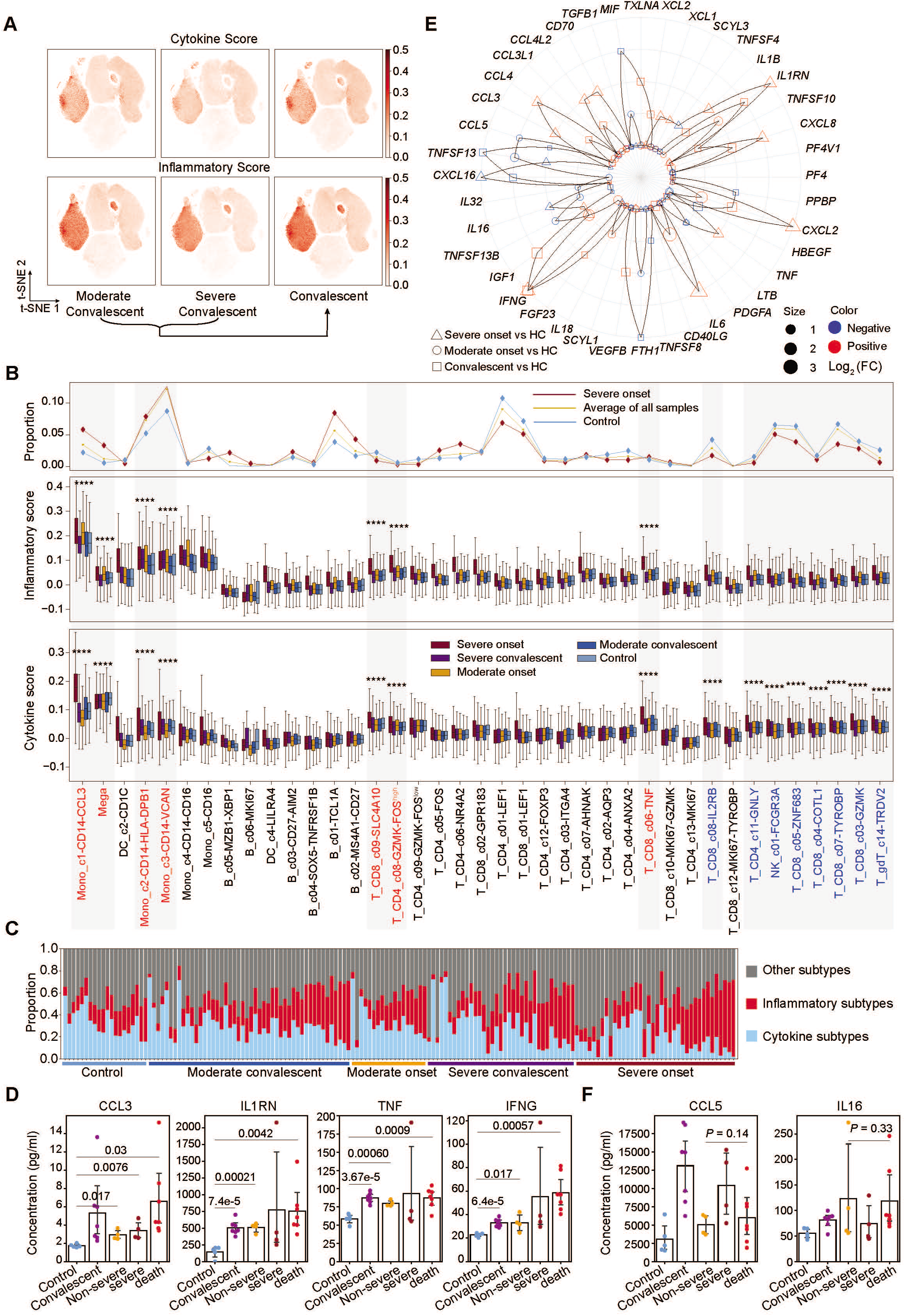
Identification of hyper-inflammatory subtypes associated with cytokine storm in PBMCs. (A) *ŕ*-SNE plots of PBMC cells colored by cytokine score (top panel) and inflammatory score (bottom panel). (B) The proportion of subtypes from healthy controls (color, n=20), severe onset (color, n=38) and average of all samples (n=159) (top panel); the inflammatory score (middle panel) and cytokine score (bottom panel) of subtypes from healthy controls (n=20), moderate convalescent (n=48), moderate onset (n=18), severe convalescent (n=35) and severe onset (n=38) patients. Significance was evaluated with Mann-Whitney rank test for each subtype versus all the other subtypes. **** *P* < 0.0001. (C) Barplots of subtypes’ (seven hyper-inflammatory cell types, eight cytokine cell types and others) frequencies across each individual samples from healthy controls (n=20), moderate convalescent (n=48), moderate onset (n=18), severe convalescent (n=35) and severe onset (n=38) patients. (D) Bar graphs showing cytokine concentration at the serum levels of CCL3, IFNG, IL1RN and TNF from healthy controls (n=5), convalescent (n=7), non-severe (n=4), severe (n=4), death case (n=7) patients. Shown are *P* values by student *t*-test. (E) The differential expression distribution of cytokines for severe onset (n=38), moderate onset (n=18) and convalescent (n=48+35) versus healthy control (n=20). The triangle represents severe onset versus healthy controls. Circle stands for moderate onset versus healthy controls. The square stands for convalescent versus healthy controls. All rings in the plot from the inside to the outside represent the range of *P* value, which are *P* > 0.05, 0.01< *P* <0.05, 0.001 < *P* <0.01 ‘ 0.0001 < *P* <0.001 and *P* < 0.0001 respectively. Red indicates positive and blue indicates negative. Size for the triangle, circle and square means log_2_ (fold change). Two-sided unpaired t test. HC, healthy control. (F) Bar graphs showing cytokine concentration at the serum levels of *CCL5* and IL16 from healthy controls (n=5), convalescent (n=7), non-severe (n=4), severe (n=4), death case (n=7) patients. Kruskal-Wallis H-test between non-severe, severe and death case. In panel (D) and panel (F), all points are shown and bars represent mean with the 95% confidence intervals. DC, dendritic cells. Mega, megakaryocytes. Mono, monocytes.

**Figure S7.**
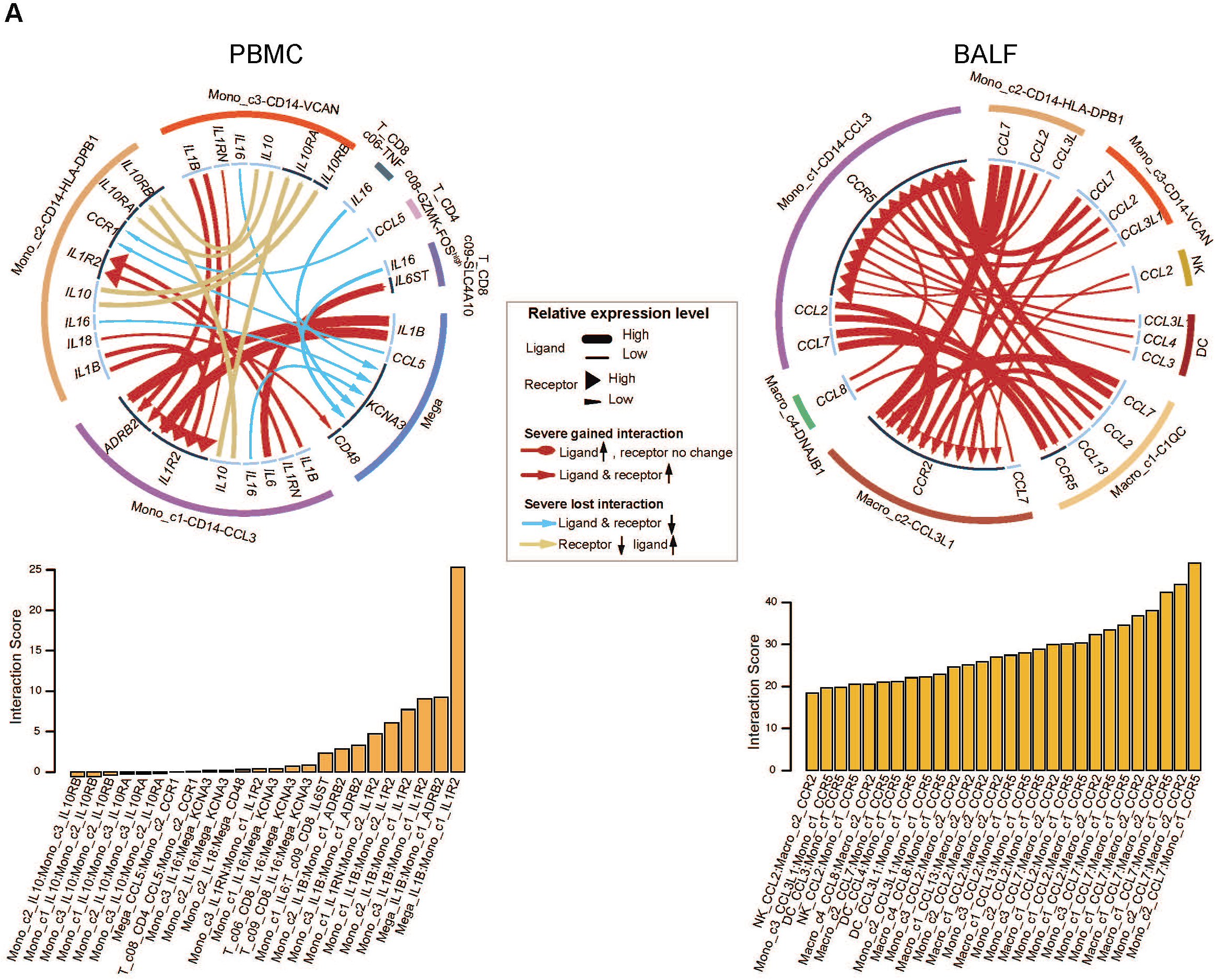
Intercellular interaction alterations among cell types between severe and moderate onset sample groups. (A). Circos plot showing the prioritized interactions mediated by ligand-receptor pairs between inflammation-related cell subtypes for each tissue, namely, PBMC (left panel) and BALF (right panel). The outer ring displays color coded cell types and the inner ring represents the involved ligand-receptor interacting pairs. The line width and arrow width are proportional to the log fold change between severe onset and moderate onset patient groups in ligand and receptor, respectively. Colors and types of lines are used to indicate different types of interactions as shown in the legend. The barplot at bottom indicates the interaction score for each ligand-receptor interaction which serves to measure the interaction strength. DC, dendritic cells. Epi, epithelial cells. Macro, macrophage cells. Mono, monocytes. Neu, neutrophils. Mega, megakaryocytes.

## Notes

### Competing Interest Statement

The authors have declared no competing interest.

## Reference

Abbasifard, M., and Khorramdelazad, H. (2020). The bio-mission of interleukin-6 in the pathogenesis of COVID-19: A brief look at potential therapeutic tactics. Life Sci 257, 118097.

Bhaskar, S., Sinha, A., Banach, M., Mittoo, S., Weissert, R., Kass, J.S., Rajagopal, S., Pai, A.R., and Kutty, S. (2020). Cytokine Storm in COVID-19-Immunopathological Mechanisms, Clinical Considerations, and Therapeutic Approaches: The REPROGRAM Consortium Position Paper. Front Immunol 11, 1648.

Bost, P., Giladi, A., Liu, Y., Bendjelal, Y., Xu, G., David, E., Blecher-Gonen, R., Cohen, M., Medaglia, C., Li, H., et al. (2020). Host-Viral Infection Maps Reveal Signatures of Severe COVID-19 Patients. Cell 181, 1475–1488 e1412.

Bray, N.L., Pimentel, H., Melsted, P., and Pachter, L. (2016). Near-optimal probabilistic RNA-seq quantification. Nat Biotechnol 34, 525–527.

Cao, X. (2020). COVID-19: immunopathology and its implications for therapy. Nat Rev Immunol 20, 269–270.

Cao, Y., Su, B., Guo, X., Sun, W., Deng, Y., Bao, L., Zhu, Q., Zhang, X., Zheng, Y., Geng, C., et al. (2020). Potent Neutralizing Antibodies against SARS-CoV-2 Identified by High-Throughput Single-Cell Sequencing of Convalescent Patients’ B Cells. Cell 182, 73–84 e16.

Catanzaro, M., Fagiani, F., Racchi, M., Corsini, E., Govoni, S., and Lanni, C. (2020). Immune response in COVID-19: addressing a pharmacological challenge by targeting pathways triggered by SARS-CoV-2. Signal Transduct Target Ther 5, 84.

Chen, Z., Ren, X., Yang, J., Dong, J., Xue, Y., Sun, L., Zhu, Y., and Jin, Q. (2017). An elaborate landscape of the human antibody repertoire against enterovirus 71 infection is revealed by phage display screening and deep sequencing. MAbs 9, 342–349.

Chua, R.L., Lukassen, S., Trump, S., Hennig, B.P., Wendisch, D., Pott, F., Debnath, O., Thurmann, L., Kurth, F., Volker, M.T., et al. (2020). COVID-19 severity correlates with airway epithelium-immune cell interactions identified by single-cell analysis. Nat Biotechnol 38, 970–979.

Costela-Ruiz, V.J., Illescas-Montes, R., Puerta-Puerta, J.M., Ruiz, C., and Melguizo-Rodriguez, L. (2020). SARS-CoV-2 infection: The role of cytokines in COVID-19 disease. Cytokine Growth Factor Rev.

Del Valle, D.M., Kim-Schulze, S., Huang, H.H., Beckmann, N.D., Nirenberg, S., Wang, B., Lavin, Y., Swartz, T.H., Madduri, D., Stock, A., et al. (2020). An inflammatory cytokine signature predicts COVID-19 severity and survival. Nat Med.

Dhama, K., Khan, S., Tiwari, R., Sircar, S., Bhat, S., Malik, Y.S., Singh, K.P., Chaicumpa, W., Bonilla-Aldana, D.K., and Rodriguez-Morales, A.J. (2020). Coronavirus Disease 2019-COVID-19. Clin Microbiol Rev 33.

Du, F., Liu, B., and Zhang, S. (2020). COVID-19: the role of excessive cytokine release and potential ACE2 down-regulation in promoting hypercoagulable state associated with severe illness. J Thromb Thrombolysis.

Gao, L., Zhou, J., Yang, S., Chen, X., Yang, Y., Li, R., Pan, Z., Zhao, J., Li, Z., Huang, Q., et al. (2020). The dichotomous and incomplete adaptive immunity in COVID-19. medRxiv, 2020.2009.2005.20187435.

Gavins, F.N., Hughes, E.L., Buss, N.A., Holloway, P.M., Getting, S.J., and Buckingham, J.C. (2012). Leukocyte recruitment in the brain in sepsis: involvement of the annexin 1-FPR2/ALX anti-inflammatory system. FASEB J 26, 4977–4989.

Giamarellos-Bourboulis, E.J., Netea, M.G., Rovina, N., Akinosoglou, K., Antoniadou, A., Antonakos, N., Damoraki, G., Gkavogianni, T., Adami, M.E., Katsaounou, P., et al. (2020). Complex Immune Dysregulation in COVID-19 Patients with Severe Respiratory Failure. Cell Host Microbe 27, 992–1000 e1003.

Gudbjartsson, D.F., Norddahl, G.L., Melsted, P., Gunnarsdottir, K., Holm, H., Eythorsson, E., Arnthorsson, A.O., Helgason, D., Bjarnadottir, K., Ingvarsson, R.F., et al. (2020). Humoral Immune Response to SARS-CoV-2 in Iceland. N Engl J Med.

Guo, C., Li, B., Ma, H., Wang, X., Cai, P., Yu, Q., Zhu, L., Jin, L., Jiang, C., Fang, J., et al. (2020a). Single-cell analysis of two severe COVID-19 patients reveals a monocyte-associated and tocilizumab-responding cytokine storm. Nat Commun 11, 3924.

Guo, Q., Zhao, Y., Li, J., Liu, J., Guo, X., Zhang, Z., Cao, L., Luo, Y., Bao, L., Wang, X., et al. (2020b). Small molecules inhibit SARS-COV-2 induced aberrant inflammation and viral replication in mice by targeting S100A8/A9-TLR4 axis. bioRxiv, 2020.2009.2009.288704.

Hadjadj, J., Yatim, N., Barnabei, L., Corneau, A., Boussier, J., Smith, N., Pere, H., Charbit, B., Bondet, V., Chenevier-Gobeaux, C., et al. (2020). Impaired type I interferon activity and inflammatory responses in severe COVID-19 patients. Science 369, 718–724.

He, J., Cai, S., Feng, H., Cai, B., Lin, L., Mai, Y., Fan, Y., Zhu, A., Huang, H., Shi, J., et al. (2020). Single-cell analysis reveals bronchoalveolar epithelial dysfunction in COVID-19 patients. Protein Cell.

Hou, Y.J., Okuda, K., Edwards, C.E., Martinez, D.R., Asakura, T., Dinnon, K.H., 3rd, Kato, T., Lee, R.E., Yount, B.L., Mascenik, T.M., et al. (2020). SARS-CoV-2 Reverse Genetics Reveals a Variable Infection Gradient in the Respiratory Tract. Cell 182, 429–446 e414.

Huang, C., Wang, Y., Li, X., Ren, L., Zhao, J., Hu, Y., Zhang, L., Fan, G., Xu, J., Gu, X., et al. (2020). Clinical features of patients infected with 2019 novel coronavirus in Wuhan, China. Lancet 395, 497–506.

Jamilloux, Y., Henry, T., Belot, A., Viel, S., Fauter, M., El Jammal, T., Walzer, T., Francois, B., and Seve, P. (2020). Should we stimulate or suppress immune responses in COVID-19? Cytokine and anti-cytokine interventions. Autoimmun Rev 19, 102567.

Jiang, N., He, J., Weinstein, J.A., Penland, L., Sasaki, S., He, X.-S., Dekker, C.L., Zheng, N.-Y., Huang, M., Sullivan, M., et al. (2013). Lineage Structure of the Human Antibody Repertoire in Response to Influenza Vaccination. Science Translational Medicine 5, 171ra119.

Jouan, Y., Guillon, A., Gonzalez, L., Perez, Y., Boisseau, C., Ehrmann, S., Ferreira, M., Daix, T., Jeannet, R., François, B., et al. (2020). Phenotypical and functional alteration of unconventional T cells in severe COVID-19 patients. Journal of Experimental Medicine 217.

Korsunsky, I., Millard, N., Fan, J., Slowikowski, K., Zhang, F., Wei, K., Baglaenko, Y., Brenner, M., Loh, P.R., and Raychaudhuri, S. (2019). Fast, sensitive and accurate integration of single-cell data with Harmony. Nat Methods 16, 1289–1296.

Kox, M., Waalders, N.J.B., Kooistra, E.J., Gerretsen, J., and Pickkers, P. (2020). Cytokine Levels in Critically Ill Patients With COVID-19 and Other Conditions. JAMA.

Kuri-Cervantes, L., Pampena, M.B., Meng, W., Rosenfeld, A.M., Ittner, C.A.G., Weisman, A.R., Agyekum, R.S., Mathew, D., Baxter, A.E., Vella, L.A., et al. (2020). Comprehensive mapping of immune perturbations associated with severe COVID-19. Sci Immunol 5.

Laouedj, M., Tardif, M.R., Gil, L., Raquil, M.A., Lachhab, A., Pelletier, M., Tessier, P.A., and Barabe, F. (2017). S100A9 induces differentiation of acute myeloid leukemia cells through TLR4. Blood 129, 1980–1990.

Lee, N., Hui, D., Wu, A., Chan, P., Cameron, P., Joynt, G.M., Ahuja, A., Yung, M.Y., Leung, C.B., To, K.F., et al. (2003). A major outbreak of severe acute respiratory syndrome in Hong Kong. N Engl J Med 348, 1986–1994.

Liao, M., Liu, Y., Yuan, J., Wen, Y., Xu, G., Zhao, J., Cheng, L., Li, J., Wang, X., Wang, F., et al. (2020). Single-cell landscape of bronchoalveolar immune cells in patients with COVID-19. Nat Med 26, 842–844.

Liberzon, A., Birger, C., Thorvaldsdottir, H., Ghandi, M., Mesirov, J.P., and Tamayo, P. (2015). The Molecular Signatures Database (MSigDB) hallmark gene set collection. Cell Syst 1, 417–425.

Lun, A.T., McCarthy, D.J., and Marioni, J.C. (2016). A step-by-step workflow for low-level analysis of single-cell RNA-seq data with Bioconductor. F1000Res 5, 2122.

Manne, B.K., Denorme, F., Middleton, E.A., Portier, I., Rowley, J.W., Stubben, C.J., Petrey, A.C., Tolley, N.D., Guo, L., Cody, M.J., et al. (2020). Platelet Gene Expression and Function in COVID-19 Patients. Blood.

Mathew, D., Giles, J.R., Baxter, A.E., Greenplate, A.R., Wu, J.E., Alanio, C., Oldridge, D.A., Kuri-Cervantes, L., Pampena, M.B., D’Andrea, K., et al. (2020a). Deep immune profiling of COVID-19 patients reveals patient heterogeneity and distinct immunotypes with implications for therapeutic interventions. bioRxiv.

Mathew, D., Giles, J.R., Baxter, A.E., Oldridge, D.A., Greenplate, A.R., Wu, J.E., Alanio, C., Kuri-Cervantes, L., Pampena, M.B., D’Andrea, K., et al. (2020b). Deep immune profiling of COVID-19 patients reveals distinct immunotypes with therapeutic implications. Science 369.

Mehta, P., McAuley, D.F., Brown, M., Sanchez, E., Tattersall, R.S., Manson, J.J., and Hlh Across Speciality Collaboration, U.K. (2020). COVID-19: consider cytokine storm syndromes and immunosuppression. Lancet 395, 1033–1034.

Melsted, P., Ntranos, V., and Pachter, L. (2019). The barcode, UMI, set format and BUStools. Bioinformatics 35, 4472–4473.

Netea, M.G., Giamarellos-Bourboulis, E.J., Dominguez-Andres, J., Curtis, N., van Crevel, R., van de Veerdonk, F.L., and Bonten, M. (2020). Trained Immunity: a Tool for Reducing Susceptibility to and the Severity of SARS-CoV-2 Infection. Cell 181, 969–977.

Ni, L., Cheng, M.-L., Zhao, H., Feng, Y., Liu, J., Ye, F., Ye, Q., Zhu, G., Li, X., Wang, P., et al. (2020a). Impaired cellular immunity to SARS-CoV-2 in severe COVID-19 patients. medRxiv, 2020.2008.2010.20171371.

Ni, L., Ye, F., Cheng, M.L., Feng, Y., Deng, Y.Q., Zhao, H., Wei, P., Ge, J., Gou, M., Li, X., et al. (2020b). Detection of SARS-CoV-2-Specific Humoral and Cellular Immunity in COVID-19 Convalescent Individuals. Immunity 52, 971–977 e973.

Nicholls, J.M., Poon, L.L., Lee, K.C., Ng, W.F., Lai, S.T., Leung, C.Y., Chu, C.M., Hui, P.K., Mak, K.L., Lim, W., et al. (2003). Lung pathology of fatal severe acute respiratory syndrome. Lancet 361, 1773–1778.

Osei-Owusu, P., Charlton, T.M., Kim, H.K., Missiakas, D., and Schneewind, O. (2019). FPR1 is the plague receptor on host immune cells. Nature 574, 57–62.

Ren, X., Zhong, G., Zhang, Q., Zhang, L., Sun, Y., and Zhang, Z. (2020). Reconstruction of cell spatial organization from single-cell RNA sequencing data based on ligand-receptor mediated self-assembly. Cell Res 30, 763–778.

Schoggins, J.W., and Rice, C.M. (2011). Interferon-stimulated genes and their antiviral effector functions. Curr Opin Virol 1, 519–525.

Schulte-Schrepping, J., Reusch, N., Paclik, D., Bassler, K., Schlickeiser, S., Zhang, B., Kramer, B., Krammer, T., Brumhard, S., Bonaguro, L., et al. (2020). Severe COVID-19 Is Marked by a Dysregulated Myeloid Cell Compartment. Cell.

Sekine, T., Perez-Potti, A., Rivera-Ballesteros, O., Strålin, K., Gorin, J.-B., Olsson, A., Llewellyn-Lacey, S., Kamal, H., Bogdanovic, G., Muschiol, S., et al. (2020). Robust T cell immunity in convalescent individuals with asymptomatic or mild COVID-19. Cell.

Silvin, A., Chapuis, N., Dunsmore, G., Goubet, A.-G., Dubuisson, A., Derosa, L., Almire, C., Hénon, C., Kosmider, O., Droin, N., et al. (2020a). Elevated calprotectin and abnormal myeloid cell subsets discriminate severe from mild COVID-19. Cell.

Silvin, A., Chapuis, N., Dunsmore, G., Goubet, A.G., Dubuisson, A., Derosa, L., Almire, C., Henon, C., Kosmider, O., Droin, N., et al. (2020b). Elevated Calprotectin and Abnormal Myeloid Cell Subsets Discriminate Severe from Mild COVID-19. Cell.

Sugimoto, M.A., Vago, J.P., Teixeira, M.M., and Sousa, L.P. (2016). Annexin A1 and the Resolution of Inflammation: Modulation of Neutrophil Recruitment, Apoptosis, and Clearance. J Immunol Res 2016, 8239258.

Sun, X., Wang, T., Cai, D., Hu, Z., Chen, J., Liao, H., Zhi, L., Wei, H., Zhang, Z., Qiu, Y., et al. (2020). Cytokine storm intervention in the early stages of COVID-19 pneumonia. Cytokine Growth Factor Rev 53, 38–42.

Sungnak, W., Huang, N., Becavin, C., Berg, M., Queen, R., Litvinukova, M., Talavera-Lopez, C., Maatz, H., Reichart, D., Sampaziotis, F., et al. (2020). SARS-CoV-2 entry factors are highly expressed in nasal epithelial cells together with innate immune genes. Nat Med 26, 681–687.

Takahashi, T., Ellingson, M.K., Wong, P., Israelow, B., Lucas, C., Klein, J., Silva, J., Mao, T., Oh, J.E., Tokuyama, M., et al. (2020). Sex differences in immune responses that underlie COVID-19 disease outcomes. Nature.

Tan, L., Wang, Q., Zhang, D., Ding, J., Huang, Q., Tang, Y.Q., Wang, Q., and Miao, H. (2020a). Lymphopenia predicts disease severity of COVID-19: a descriptive and predictive study. Signal Transduct Target Ther 5, 33.

Tan, W., Lu, Y., Zhang, J., Wang, J., Dan, Y., Tan, Z., He, X., Qian, C., Sun, Q., Hu, Q., et al. (2020b). Viral Kinetics and Antibody Responses in Patients with COVID-19. medRxiv, 2020.2003.2024.20042382.

Traag, V.A., Waltman, L., and van Eck, N.J. (2019). From Louvain to Leiden: guaranteeing well-connected communities. Sci Rep 9, 5233.

Vabret, N., Britton, G.J., Gruber, C., Hegde, S., Kim, J., Kuksin, M., Levantovsky, R., Malle, L., Moreira, A., Park, M.D., et al. (2020). Immunology of COVID-19: Current State of the Science. Immunity 52, 910–941.

Vogl, T., Tenbrock, K., Ludwig, S., Leukert, N., Ehrhardt, C., van Zoelen, M.A.D., Nacken, W., Foell, D., van der Poll, T., Sorg, C., et al. (2007). Mrp8 and Mrp14 are endogenous activators of Toll-like receptor 4, promoting lethal, endotoxin-induced shock. Nature Medicine 13, 1042–1049.

Wauters, E., Van Mol, P., Garg, A.D., Jansen, S., Van Herck, Y., Vanderbeke, L., Bassez, A., Boeckx, B., Malengier-Devlies, B., Timmerman, A., et al. (2020). Discriminating Mild from Critical COVID-19 by Innate and Adaptive Immune Single-cell Profiling of Bronchoalveolar Lavages. bioRxiv, 2020.2007.2009.196519.

Wen, W., Su, W., Tang, H., Le, W., Zhang, X., Zheng, Y., Liu, X., Xie, L., Li, J., Ye, J., et al. (2020). Immune cell profiling of COVID-19 patients in the recovery stage by single-cell sequencing. Cell Discov 6, 31.

Wilk, A.J., Rustagi, A., Zhao, N.Q., Roque, J., Martinez-Colon, G.J., McKechnie, J.L., Ivison, G.T., Ranganath, T., Vergara, R., Hollis, T., et al. (2020). A single-cell atlas of the peripheral immune response in patients with severe COVID-19. Nat Med 26, 1070–1076.

Wolf, F.A., Angerer, P., and Theis, F.J. (2018). SCANPY: large-scale single-cell gene expression data analysis. Genome Biol 19, 15.

Wolock, S.L., Lopez, R., and Klein, A.M. (2019). Scrublet: Computational Identification of Cell Doublets in Single-Cell Transcriptomic Data. Cell Syst 8, 281–291 e289.

Xie, X., Liu, M., Zhang, Y., Wang, B., Zhu, C., Wang, C., Li, Q., Huo, Y., Guo, J., Xu, C., et al. (2020). Single-cell Transcriptomic Landscape of Human Blood Cells. Natl Sci Rev.

Yan, X., Li, F., Wang, X., Yan, J., Zhu, F., Tang, S., Deng, Y., Wang, H., Chen, R., Yu, Z., et al. (2020). Neutrophil to lymphocyte ratio as prognostic and predictive factor in patients with coronavirus disease 2019: A retrospective cross-sectional study. J Med Virol.

Yang, Y., Shen, C., Li, J., Yuan, J., Wei, J., Huang, F., Wang, F., Li, G., Li, Y., Xing, L., et al. (2020). Plasma IP-10 and MCP-3 levels are highly associated with disease severity and predict the progression of COVID-19. J Allergy Clin Immunol 146, 119–127 e114.

Yu, G., Wang, L.G., Han, Y., and He, Q.Y. (2012). clusterProfiler: an R package for comparing biological themes among gene clusters. OMICS 16, 284–287.

Yu, K., He, J., Wu, Y., Xie, B., Liu, X., Wei, B., Zhou, H., Lin, B., Zuo, Z., Wen, W., et al. (2020). Dysregulated adaptive immune response contributes to severe COVID-19. Cell Res 30, 814–816.

Zhang, C., Gadue, P., Scott, E., Atchison, M., and Poncz, M. (1997). Activation of the megakaryocyte-specific gene platelet basic protein (PBP) by the Ets family factor PU.1. J Biol Chem 272, 26236–26246.

Zhang, F., Gan, R., Zhen, Z., Hu, X., Li, X., Zhou, F., Liu, Y., Chen, C., Xie, S., Zhang, B., et al. (2020a). Adaptive immune responses to SARS-CoV-2 infection in severe versus mild individuals. Signal Transduct Target Ther 5, 156–156.

Zhang, J., Sze, D.M., Yung, B.Y., Tang, P., Chen, W.J., Chan, K.H., and Leung, P.H. (2016). Distinct expression of interferon-induced protein with tetratricopeptide repeats (IFIT) 1/2/3 and other antiviral genes between subsets of dendritic cells induced by dengue virus 2 infection. Immunology 148, 363–376.

Zhang, J.Y., Wang, X.M., Xing, X., Xu, Z., Zhang, C., Song, J.W., Fan, X., Xia, P., Fu, J.L., Wang, S.Y., et al. (2020b). Single-cell landscape of immunological responses in patients with COVID-19. Nat Immunol 21, 1107–1118.

Zhang, L., Pang, R., Xue, X., Bao, J., Ye, S., Dai, Y., Zheng, Y., Fu, Q., Hu, Z., and Yi, Y. (2020c). Anti-SARS-CoV-2 virus antibody levels in convalescent plasma of six donors who have recovered from COVID-19. Aging (Albany NY) 12, 6536–6542.

Zhang, L., Yu, X., Zheng, L., Zhang, Y., Li, Y., Fang, Q., Gao, R., Kang, B., Zhang, Q., Huang, J.Y., et al. (2018). Lineage tracking reveals dynamic relationships of T cells in colorectal cancer. Nature 564, 268–272.

Zhou, C., Gao, C., Xie, Y., and Xu, M. (2020a). COVID-19 with spontaneous pneumomediastinum. Lancet Infect Dis 20, 510.

Zhou, Y., Fu, B., Zheng, X., Wang, D., Zhao, C., qi, Y., Sun, R., Tian, Z., Xu, X., and Wei, H. (2020b). Pathogenic T cells and inflammatory monocytes incite inflammatory storm in severe COVID-19 patients. Natl Sci Rev, nwaa041.

